# A rhizobial extracellular vesicle-conveyed auxin transporter mediates phytohormone mobilization and optimizes rhizobium-legume symbiosis

**DOI:** 10.64898/2026.07.10.737661

**Authors:** Natalia Moreno-de Castro, Mustafa Safa Karagöz, Irene Herrero Gómez, Paula Ayala-García, Susanne Sievers, Mathias Müsken, Joaquín Giner-Lamia, Irene Jiménez-Guerrero, Francisco Pérez-Montaño, José Manuel Borrero-de Acuña

## Abstract

Bacterial extracellular vesicles (bEVs) are emerging as key players in interkingdom communication, yet their role in delivering functional proteins to host cells during symbiosis remains unexplored. This study shows that *Sinorhizobium fredii* HH103 packages a PIN-like auxin transporter, AuxT, into bEVs that traffic within the peribacteroid space of soybean nodules. AuxT is chromosomally encoded and constitutively expressed, genetically uncoupled from the flavonoid-inducible auxin biosynthesis machinery located on the symbiotic plasmid. Structural prediction reveals that AuxT adopts an eight-transmembrane-helix architecture with striking homology to plant PIN auxin exporters, despite negligible sequence identity. Molecular docking demonstrates that AuxT binds indole-3-acetic acid within a central cavity, with dimerization inducing ligand-specific conformational changes consistent with transport activity. The *auxT* mutant exhibits significant symbiotic defects including reduced shoot biomass, nodule number, and nodule mass that are fully restored by complementation. Critically, AuxT-enriched bEVs contain elevated auxin levels, and nodules colonized by the complemented strain accumulate more auxin specifically within the bEVs peribacteroid space, while bacteroids themselves show no auxin retention. We postulate that bEV-associated AuxT mediates localized auxin export into the symbiosome, modulating the host hormonal environment to optimize symbiotic development. This work reveals a previously unrecognized mechanism of interdomain hormonal modulation, where a bacterium uses a structurally convergent transporter and vesicular delivery to actively shape host physiology and to improve the symbiotic performance.

## Introduction

Bacterial extracellular vesicles (generically abbreviated as bEVs) are spherical, nano-sized structures (typically 20-400 nm) with either a single or double lipid bilayer, which are released into the surrounding environment by organisms across all domains of life (Gill *et al*., 2019; Toyofuku *et al*., 2023). In Gram-negative bacteria, bEVs originate through two primary mechanisms: explosive cell lysis and membrane blebbing. Explosive cell lysis generates E-type outer membrane vesicles (E-OMVs) or outer-inner membrane vesicles (E-OIMVs), whereas blebbing of the outer membrane alone or both the outer and inner membranes promotes the formation of B-type OMVs (B-OMVs) or OIMVs (B-OIMVs) (Toyofuku *et al*., 2019; Turnbull *et al*., 2016). Subsequently, depending on the biogenesis mechanism, the cargo of each bEV type differs. Owing to their selectively packaged cargo, which differs from that of other cellular compartments, bEVs are regarded as an ancient intercellular communication pathway, conceptually classified as the Type 0 secretion system (T0SS) (Guerrero-Mandujano *et al*., 2017). Unlike classical secretion systems that release molecules into the extracellular space or directly translocate effectors or DNA into target cells, bEVs export a diverse range of cargo, including lipids, hydrophobic molecules, insoluble materials, and virulence factors (Biller *et al*., 2021; Villageliu & Samuelson, 2022). In contrast to other secretion systems, bEV-mediated transport provides several advantages for the cargo, including the synergistic activity of molecular cocktails, protection against degradation, targeted delivery, and maintenance of critical concentrations for bioactivity (Bitto *et al*., 2023; Renelli *et al*., 2004; Wang *et al*., 2023). Indeed, cargo compounds are concentrated within bEVs, ensuring the delivery of a biologically relevant dose to target cells, a phenomenon known as quantal secretion (Toyofuku *et al*., 2017). bEV-mediated biological processes that impact other organisms encompass bacterial killing, DNA transfer, effector delivery, immunomodulation, and the delivery of bioactive compounds (Fulsundar *et al*., 2014; Kadurugamuwa & Beveridge, 1996; Li *et al*., 2022). These latter functions are particularly important for plant pathogenic and mutualistic bacteria, including rhizobia.

Rhizobia are soilborne α- and ß-proteobacteria capable of establishing symbiotic relationships with legumes. Within specialized root organs called nodules, rhizobia convert atmospheric nitrogen into ammonia, increasing nitrogen bioavailability for the host plant and thereby enhancing crop yields (Oldroyd, 2013; Roy *et al*., 2020). The onset of a complex dialogue culminating with the rhizobium-legume symbiosis is ignited by the exudation of flavonoids from legume roots to the rhizosphere. Specific flavonoids are recognized by the transcriptional regulator NodD, resulting in the activation of the expression of nodulation (*nod*) genes (Fisher & Long, 1989; Peck *et al*., 2006; Wassem *et al*., 2017). These genes encode proteins involved in the synthesis and export of compatible lipo-chito-oligosaccharide molecules, also known as Nod factors (NFs). In turn, these molecules are specifically recognized by plant LysM receptor-like kinases, deploying a downstream signaling cascade that includes the upregulation of various symbiotic plant genes (Luo *et al*., 2023; Masson-Boivin *et al*., 2009; Zipfel & Oldroyd, 2017). This signaling process leads to rhizobium-specific intracellular colonization, typically via the formation of infection threads within root hairs, culminating in nodule development and infection (Oldroyd *et al*., 2011). Interestingly, recent studies have underscored the importance of bEVs at both early and late stages of the symbiotic process, highlighting increased complexity driven by an expanded and tightly intertwined network of molecular mechanisms underlying this intimate relationship. Notably, bEVs derived from *Rhizobium etli* and *Sinorhizobium fredii* that had been exposed to *nod* gene-inducing flavonoids were shown to deform the root hairs of host legumes, indicating the presence of biologically active NFs within these molecular conveyors (Li *et al*., 2022; Taboada *et al*., 2019). In particular, this observation has been confirmed for bEVs from *S. fredii* HH103, which selectively package and transport high-molecular-weight NFs, delivering a concentrated, biologically active dose that promotes nodule primordia formation and enhances overall nodulation across different host legumes (Moreno-de Castro *et al*., 2026). During the late stages of the symbiotic process intense bEV trafficking also occurs within the peribacteroid space of nodules, which constitutes the interface between the host plant membrane and the nitrogen-fixing bacteroids. In our previous work, we isolated bEVs from this space of several symbiotic pairs demonstrating that they carry proteins potentially required for sustaining symbiotic function and bacteroid viability (Ayala-García al., 2024a). Among them, we found a putative auxin efflux carrier (CCE98274.1), that structurally resembles the plant-borne PIN8 auxin transporter. This protein was identified as a constituent of the bEVs released by *S. fredii* HH103 bacteroids within host-legume nodules, potentially aiding in the uptake of this phytohormone by the host plant cells. Plant PIN8 transporters, which are non-canonical PINs primarily localized on organellar membranes such as the endoplasmic reticulum (ER), function as constitutively active auxin efflux carriers that operate independently of activating kinases (Ding *et al*., 2012; Mravec *et al*., 2009). Structural and biophysical analyses of PIN8 demonstrated that it employs an elevator-type transport mechanism, with movement driven by the negative charge of the auxin anion, supporting a uniport mechanism independent of obligatory proton or ion gradients (Ung *et al*., 2022). In plants, PIN8 regulates auxin concentration by functioning as a constitutively active efflux carrier that exports auxin from the cytosol into the ER lumen. This sequestration effectively reduces the pool of cytosolic and nuclear auxin, thereby negatively modulating auxin-dependent gene expression responses (Barbez & Kleine-Vehn, 2013; Weijers & Wagner, 2016, Han *et al*., 2021). These transporters, belonging to the bile/arsenite/riboflavin transporter (BART) superfamily, are considered plant-specific and display dynamic subcellular localization in response to developmental and environmental cues (Friml, 2022).

Auxin is a crucial phytohormone regulating almost all aspects of plant growth and development, playing a dynamic role in coordinating the legume-rhizobia symbiosis by promoting both the initial epidermal infection and subsequent root nodule organogenesis (Velandia *et al*., 2022). Auxins are endogenously synthesized by plants through both tryptophan-dependent and tryptophan-independent pathways, ensuring local and systemic hormone availability during development (Solanki & Shukla, 2023). However, endogenous production does not eliminate the need for transport systems, as the biological activity of auxin largely depends on its directional and tightly regulated distribution across tissues and subcellular compartments, a process known as polar auxin transport (PAT) (Friml, 2022; Han *et al*., 2021). During symbiosis, auxin functions as a positive regulator of epidermal infection, where its’ signaling is activated in root hairs following NF treatment to facilitate infection thread formation (Breakspear *et al*., 2014; Nadzieja *et al*., 2018; Velandia *et al*., 2022). This hormone is also indispensable for cortical nodule organogenesis, requiring a local accumulation in dividing cortical cells to initiate and maintain the primordium (Luo *et al*., 2023; Suzaki *et al*., 2015).

Interestingly, soil-borne bacteria in general and rhizobia in particular can produce auxins constitutively and/or in response to flavonoids, the legume-secreted signals, because in this last case the synthesis of this critical phytohormone is integrated into the NodD regulatory cascade, mirroring the host-specific regulation of *nod* genes (Theunis *et al*., 2004; Tullio *et al*., 2019). Knockout mutations in the bacterial auxin synthesis genes have revealed complex symbiotic phenotypes across rhizobial strains. In *S. fredii* NGR234, mutation of *y4wE* resulted in a significant reduction of free auxins found within nodules formed on *Vigna unguiculata* and *Tephrosia vogelii* (Theunis *et al*., 2004). However, the establishment of symbiosis was not abolished, as no significant differences were found in the symbiotic parameters analyzed in comparison to those induced by the wild-type strain (Theunis *et al*., 2004). Conversely, in *Rhizobium tropici* CIAT 899, mutant defective in *y4wF* auxin synthesis gene showed a delay in nodulation kinetics *on Phaseolus vulgaris* (Tullio *et al*., 2019). Another distinction between both strains is the substantial constitutive production observed in NGR234, whereas in CIAT 899, most pathways remain dormant or exhibit basal activity in the absence of flavonoid-induction (Theunis *et al*., 2004; Tullio *et al*., 2019). Notably, although rhizobia and other soil bacteria can synthesize auxins, current knowledge is limited to bacterial auxin biosynthesis, delivery pathways, and their contribution to symbiotic phenotypes. To date, there are no reports of a PIN-like auxin efflux transporter packaged within rhizobial bEV cargo to facilitate the vesicle-mediated delivery of this phytohormone to host cells.

In this study, we report for the first time that *S. fredii* HH103 encodes a PIN-like auxin transporter, AuxT, which facilitates auxin translocation into the periplasm and subsequent loading into bEVs. We show that AuxT is selectively co-packaged with auxins into bEVs and co-delivered to the symbiosome interface. Through integrated genomic, structural, biochemical, microscopy, and symbiotic analyses, we demonstrate that AuxT is required for optimal symbiosis and enhances auxin loading into bEVs under both free-living and symbiotic conditions. This work reveals a previously unrecognized mechanism of interdomain hormonal modulation that is ultimately crucial for efficient plant growth promotion mediated by rhizobia.

## Materials and methods

### Growth conditions and construction of plasmids and bacterial strains

Bacterial strains used in this work are listed in **Table S1**. Rhizobial strains were grown at 28°C on tryptone yeast (TY) medium (Beringer, 1974), mannitol minimal (MM) medium (Robertsen *et al*., 1981) with different mannitol concentrations (10 g mL^-1^ for MM and 3 g mL^-1^ for MM3) or yeast-extract mannitol (YM) medium (Vincent, 1970b). *E. coli* strains were cultured on Luria-Bertani (LB) medium (Sambrook *et al*., 1989) at 37°C. In the case of liquid media, cultures were grown with continuous shaking at 180 r.p.m. When required, media were supplemented with appropriate antibiotics at the concentrations described in Lamrabet *et al*. (1999). Antibiotics were used at the following concentrations (µg mL^-1^): rifampicin (Rif) 50 for *S. fredii* and kanamycin (Km) 50 for *S. fredii* and 25 for *E. coli.* Genistein (for *S. fredii* HH103) and apigenin (for *R. tropici* CIAT 899), which are nod gene-inducing flavonoids, were dissolved in absolute ethanol at a concentration of 1 µg mL^−1^ to achieve a final concentration of 3.7 µM. 5-aminolevulinic acid (ALA), a nutrient required for *E. coli* ST18 to grow, was dissolved in water at a concentration of 50 mg mL^−1^ to obtain a final concentration of 50 µg mL^-1^ (Thoma & Schobert, 2009).

Primers and plasmids used in this work are listed in **tables S2 and S3**, respectively. The *auxT* knockout mutant (Δ*auxT*) was generated by deletion mutagenesis. A 5’ flanking region of *auxT* was amplified using primers auxt_EcoRI_Fw_A (P1) and auxt_Rev_B (P2), while a 3’ flanking region of *auxT* was amplified using primers auxt_Fw_C (P3) and auxt_BamHI_Rev_D (P4). Subsequently, the two flanking regions were fused by overlap PCR using primers P1 and P4, generating an *auxT* version containing an internal deletion. The resulting 1129 bp PCR product was cloned into the vector pk18*mobsacB* (Schäfer *et al*., 1994), by previous digestion with *Eco*RI and *Bam*HI enzymes of both the fragment and the plasmid and sequent ligation with the enzyme T4 DNA ligase. The pK18*mobsacB* vector is a non-replicative vector (suicide plasmid) in rhizobia and confers kanamycin resistance (Km^R^, 50 ug mL^-1^). The resulting plasmid was first transformed into *E. coli* DH5α strain (hereafter DH5α), isolated via Plasmid Miniprep System and transformed into *E. coli* ST18 strain (hereafter ST18). This plasmid was subsequently transferred from ST18 to HH103 by biparental mating, as described by (Simon, 1984). Single recombinants were selected on TY Km medium. Next, single recombinants were grown in liquid TY medium at 28 °C for approximately 5 days to allow spontaneous double recombination. Cultures were then plated onto TY Sac medium to select double recombinants, as cells carrying the pk18*mobsacB* plasmid are unable to grow in the presence of sucrose. Putative double recombinants were further screened by replica plating onto TY Sac, TY Km and LB plates. Colonies that grew exclusively on Ty Sac plates were selected as double recombinants. Mutation of *auxT* was confirmed by PCR amplification and DNA sequencing using the primers auxt_ext_Fw (P13) and auxt_ext_Rev (p14).

To obtain the complemented *auxT* mutant strain (Δ*auxT*-_p_*auxT*), primer pairs Pauxt_EcoRI_Fw (P5) and auxt_stpII_BamHI_Rev (P6) were used for amplifying the fragment containing *auxT* preceded 500 bp of its native upstream regulatory region, corresponding to its putative natural promoter, and followed by a strep-tag II. The resulting 1731 bp PCR product was then cloned into the vector pSEVA 221 (Silva-Rocha *et al*., 2013b), by digestion and ligation as previous described. This vector confers resistance to kanamycin (Km^R^ 50 ug mL^-1^). The resulting plasmid was transformed and transferred as previously described.

The *m*Cherry*-auxT* strain was generated by introducing the construct pSEVA221::*P_auxT_-mCherry::auxT* into the parental *S. fredii* HH103 strain. To assemble this construct, the native *auxT* promoter region (P*_auxT_*) was amplified using primers Pauxt_EcoRI_Fw (P7) and Pauxt_Rev (P8), incorporating an *Eco*RI restriction site at the 5’ end. The *mCherry* coding sequence was amplified using primer mcherry_Fw (P9) and mcherry_FL_Rev (P10), while the *auxT* gene was amplified using primers auxt_FL_Fw (P11) and auxt_BamHI_Rev (P12), introducing a *Bam*HI restriction site at the 3’ end. Primers P10 and P11 were designed to encode a flexible linker (FL), allowing the insertion of this linker between mCherry and AuxT, to make the proteins more likely to be functional.

Primers P8, P9, P10 and P11 contained overlapping sequences that enabled the fusion of the three fragments by overlap extension PCR. The fragments were assembled in the following order: an overhang corresponding to the flanking regions that had to be fused, so the three fragments were fused in this order: P*_auxT_*, *mCherry* and *auxT* by overlap PCR using primers P7 and P12. The resulting 2437 bp PCR product was cloned into the pSEVA 221 vector by digestion and ligation as previously described. The fragments were assembled in the following order: P*_auxT_*, *mCherry* and *auxT*, using primers P7 and P12. The resulting 2437 bp PCR product was digested with *Eco*RI and *Bam*HI and ligated into the pSEVA221 vector, as described above. The recombinant plasmid was subsequently introduced into *S. fredii* HH103.

### In silico analyses

To assess structural similarity between AuxT and other proteins, a structure-based homology search was performed using Foldseek (https://search.foldseek.com/search). Foldseek is a fast and sensitive protein structure search and alignment tool that enables the comparison of experimental and predicted protein structures (van Kempen *et al*., 2024), leveraging structural models available in multiple databases, including the AlphaFold Protein Structure Database (AFDB) (Varadi *et al*., 2022; Varadi *et al*., 2024). The predicted structure of AuxT was queried against the AFDB to identify proteins exhibiting high structural homology. To compare the amino acid sequence of AuxT with those of the proteins identified in the Foldseek search, pairwise sequence alignments were performed using the BLAStp (Boratyn *et al*., 2013) tool from the NCBI platform. In addition, to identify sequence-based homologs of AuxT, a BLASTp search was conducted against the NCBI non-redundant protein database (nr) using default parameters. *S. fredii* HH103 AuxT homologues (*SFHH103_01028, SFHH103_01341* and *SFHH103_03783*) were aligned using T-Coffee (Di Tomaso *et al*., 2011). Genomic context analysis of the *auxT* homologues within the *Sinorhizobium/Ensifer* clade was conducted using GeCoViz (Botas *et al*., 2022). The presence of an N-terminal signal peptide in AuxT was assessed using SignalP v6.0, a tool that predicts signal peptides and their cleavage sites in proteins (Teufel *et al*., 2022). The transmembrane topology of AuxT was analyzed using DeepTMHMM, which predicts the presence and arrangement of transmembrane helices (Hallgren *et al*., 2022). The subcellular localization of AuxT was predicted using PSORTb v3.0, a tool designed to predict the localization of proteins (Yu *et al*., 2010).

To elucidate the three-dimensional structure of AuxT, structural models of the monomeric, dimeric and tetrameric forms were generated using the RoseTTaFold All-Atom (RFAA) platform (Krishna *et al*., 2024). For structural alignment between AuxT and its structural homologs, three-dimensional models were retrieved from AlphaFold3 (Jumper *et al*., 2021), and structural superimpositions were performed using PyMOL (C., 2015), where automated scripts were utilized for visualization and the calculation of root mean square deviation (RMSD) values to quantify structural similarity.

For molecular docking analysis, a computational simulation technique used to predict the most probable three-dimensional orientation adopted by a macromolecule upon binding to a small molecule, protein structures were modelled using the RFAA platform. The resulting structural models were subsequently used as input for docking simulations performed in the RFAA platform.

### Field Emission Scanning Electron Microscopy (FESEM) of bacterial cultures

*S. fredii* HH103 was inoculated in MM liquid medium from a fresh plate and incubated with the conditions previously described, till an optical density at 600 nm (OD_600_) of 0.8-1 was reached (≈24 h). Afterwards, samples were fixed with 2% glutaraldehyde, left for 30 min and then formaldehyde was added to a final concentration of 5%. After 1 h on ice or standing overnight at 7 °C fixed bacteria were washed twice for 10 min with TE buffer (20 mM TRIS, 1 mM EDTA, pH 6.9), dehydrated with a graded series of acetone (10, 30, 50, 70, 90%) for 15 min for each step on ice. Samples in the 100% acetone step were allowed to reach room temperature before another change in 100% acetone. Samples were then subjected to critical-point drying with liquid CO_2_. Dried samples were fixed onto aluminium stubs with a conductive adhesive tape and covered with a gold/palladium film by sputter coating before examination in a field emission scanning electron microscope Zeiss LIBRA 120 (Zeiss, Oberkochen) using the Everhart Thornley HESE2 detector and the inlens SE detector in a 25:75 ratio with an acceleration voltage of 5 kV using the SEM software 5.5.

### Isolation of rhizobial EVs

Isolation of bEVs from free living conditions was done following the steps described by (Ayala-García *et al*., 2024c), with some modifications. The different strains were inoculated in TY liquid medium and incubated for 72 h with the conditions previously described. Subsequently, these saturated cultures were inoculated in a flask containing 200 mL of MM medium with an initial OD_600_ of 0.1 and further incubated for 24 h under the same conditions as previously described. Next, the cultures were centrifuged at 5,000 x g for 20 minutes at 4 °C, the pellet was discarded and the supernatants were centrifuged at 7,000 x g for 20 minutes at 4 °C. The supernatants were filtered with a 0.2 μm polyethersulfone bottle top filter, and the eluent concentrated from 200 mL to 25 mL with the Vivaflow 50R easy load ultrafiltration system using Vivaflow 50R 30,000 (kDa) MWCO Hydrosart membranes and following manufacturer instructions. Lastly, the concentrated supernatants were filtered with a 0.2 μm filter and transferred to 30 mL ultracentrifuge tubes and ultracentrifuged, using a Himac P70AT-1759 Rotor (Ultracentrifuge CP 90NX, Eppendorf Himac Technologies, Japan), at 150,000 x g at 4 °C for 2 h. After centrifugation, the supernatants were discarded and the pellets (containing the bEVs) resuspended in 100 µL of ice-cold HEPES buffer.

Isolation of bEVs from bacteroids was performed following the protocol described by (Ayala-García *et al*., 2024c), with modifications. Nodules were ground using a sterile mortar and pestle in MMS buffer [40 mM 3-(4-morpholino)-propanesulfonic acid, 20 mM KOH, 2 mM MgSO_4_, 0.3 M sucrose, pH 7.0]. Debris derived from plant tissues was removed by collecting the liquid from the mortar and filtering it with a 40 µm cell strainer cap (BD Falcon, BD Biosciences, USA). The eluents were centrifuged at 0.1 x g for 5 minutes and the resulting pellet discarded, then the supernatants were centrifuged at 2.2 x g for 5 minutes and the resulting pellets, which correspond to the bacteroids fraction, discarded or kept, depending on the experiment. Next, the supernatants were centrifuged at 7,000 x g and 4 °C for 20 minutes. Lastly, the supernatants were filtrated using 0.2 µm filter and the filtered supernatants were transferred to 30 mL ultracentrifuge tubes and ultracentrifuged as previously described. After this centrifugation, the supernatants were discarded and the pellets (containing bEVs) resuspended in 100 µL of ice-cold HEPES buffer.

### bEV integrity assessment and quantification

Integrity and quantification assessment of purified bEVs was performed following a protocol described in (Ayala-García *et al*., 2024c). For transmission electron microscopy (TEM), 30 - 50 droplets of the sample were deposited onto a fine carbon film to promote vesicle adhesion. After 1 min of incubation, a 300-mesh copper grid was employed to lift the carbon film. The grids were then washed twice with droplets of distilled water and subsequently stained with 4% (w/v) uranyl acetate. After 1 min, excess stain was carefully removed with filter paper, and the grids were allowed to dry under a 60 W light bulb. Samples were examined using a Libra 120 transmission electron microscope (Zeiss, Oberkochen, Germany) operated at an accelerating voltage of 120 kV and calibrated magnifications. Image contrast and brightness adjustments, as well as particle size measurements, were performed using WinTEM software v01.06. bEV yields were quantified by scattering-light-reliant nanoparticle tracking analysis using the NanoSight NS300 nanoparticle analyzer (Malvern Panalytical, United Kingdom), applying a monochromatic laser beam at 488 nm and taking 90 s videos at 23 °C. To quantify and size bEVs, each video was analyzed with the nanoparticle tracking analysis software following the manufacturer’s instructions.

### Detection of AuxT transporter in the bEV fractions by immunoblot

The presence of AuxT in bEVs derived from bacteroids and free-living conditions was evaluated by Western blot analysis using the Strep-Tactin HRP detection system (IBA Lifesciences, Germany). bEVs aliquots were in SDS-PAGE sample loading buffer to a final composition of 62.5 mM Tris–HCl (pH 6.8), 2% (w/v) SDS, 10% (v/v) glycerol, 5% (w/v) β-mercaptoethanol, and 0.001% (w/v) bromophenol blue. Samples were loaded onto stain-free polyacrylamide gels (Bio-Rad, USA) and separated by electrophoresis. Proteins were subsequently transferred to nitrocellulose membranes (Bio-Rad, USA).

For protein transfer, gel and membrane were assembled in direct contact and sandwiched between Whatman filter papers pre-soaked in transfer buffer [48 mM Tris, 39 mM glycine, 1.3 mM SDS, and 20% methanol]. The transfer assembly was placed in a Mini Trans-Blot electrophoretic transfer cell (Bio-Rad, USA) and run at 25 V and 1 A for 15 min. Following transfer, the membrane was blocked overnight at 4 °C with gentle agitation in PBS blocking buffer [PBS (137 mM NaCl, 2.7 mM KCl, 10 mM phosphate buffer), 3% BSA, and 0.05% (v/v) Tween 20] to prevent non-specific antibody binding. The membrane was then washed three times for 5 min each with PBS-Tween buffer [PBS containing 0.1% (v/v) Tween 20] at room temperature. Subsequently, 10 mL of PBS-Tween buffer and 10 µL of Strep-Tactin HRP (previously pre-diluted 1:100 in Enzyme dilution buffer [PBS buffer, 0.2% BSA, 0.1% Tween]) was added to the membrane, that was incubated overnight at 4 °C with gentle shaking. After incubation, the membrane was washed three times with PBS-Tween buffer for 5 minutes each. Protein bands were visualized by applying 250 µL of the chemiluminescent substrate Immobilon Forte HRP (Merck KGaA, Germany). Chemiluminescent signals were detected using the LI-COR Odyssey FC 700/800/Chemi imaging system and analyzed with Image Studio Lite software (LI-COR Biosciences, USA).

### Evaluation of AuxT occurrence in the symbiosome

To assess the presence of AuxT in the symbiosome, an *in vivo* analysis of *auxT* gene expression in *L. burttii* nodules were performed by confocal fluorescence microscopy, following the protocol described by (Kelly *et al*., 2018) with minor modifications. *L. burttii* seeds were chemically scarified surface-sterilized and germinated as previously described by (Birgitte Heckmann *et al*., 2011). Germinated seeds were transferred to 12 × 12 cm Petri dishes containing 50 mL of ByD medium (Kelly *et al*., 2018), with two-thirds of the surface solidified at an incline and a sterile filter paper moistened with sterile distilled water. Between seven and ten seeds were placed per plate. Plates were sealed with surgical tape and incubated at an appropriate inclination to maintain seedlings in a vertical position for 6 days with a 16 h photoperiod at 21 °C in the light and 18 °C in the dark.

After this period, seedlings were inoculated with *S. fredii* HH103 strains carrying either the empty vector pSEVA221 (HH103-_p_221) or pSEVA221::P*_auxT_*-*mCherry::auxT* (HH103 mCherry-AuxT), which synthesizes the AuxT protein tagged with an N-terminal *m*Cherry protein. Bacterial cultures were inoculated at an optical density of OD□□□ = 0.6. Inoculated seedlings were grown under the same conditions until the formation of nitrogen-fixing nodules. For confocal microscopy analysis, nodules approximately 30 days post-inoculation were collected from inoculated plants to assess nodule occupancy by rhizobia.

For confocal microscopy sample preparation, nodules were embedded in 6% (w/v) agarose dissolved in distilled water and sectioned into 50 µm slices using a Leica VT1000S vibratome (Wetzlar, Germany). Sections were stained with 0.04% (w/v) calcofluor white to visualize nodule cell walls, following the protocol described by Kawaharada *et al*. (2017), and examined using a Leica Stellaris 8 SPE confocal fluorescence microscope (Leica Microsystems, Jena, Germany).

### Auxin quantification in rhizobia

Quantification of auxin production by *S. fredii* HH103 and *R. tropici* CIAT 899 was carried out by using Salkowski colorimetric assay (Glickmann & Dessaux, 1995), as described previously by (Fierro-Coronado *et al*., 2014). *S. fredii* HH103 and *R. tropici* CIAT 899 were inoculated in 20 mL of TY medium with tryptophan (500 mg L^−1^), supplemented with 20 µL of the inducing flavonoid (genistein for HH103 and apigenin for CIAT 899) or EtOH 100% (control) and incubated for 96 h at 28°C, with continuous shaking of 180 r.p.m. Afterwards, 1 mL of each culture was centrifuged at 12,000 x g speed for 1 minute to remove cells. Next, the supernatant was distributed in wells of a 96 well plate, adding 40 µL of the supernatant to each well. 160 µL of Salkowski reagent [H_2_SO_4_ 2.65 M and FeCl_3_ 9.15 mM in H_2_O] were added to the wells containing the supernatants. The mix was incubated for 20 minutes. To quantify the auxin production the absorbance was measured at 535 nm in a plate reader. Every experiment was performed in three biological replicates with eight technical replicates each time.

For auxin quantification in bEVs, vesicles derived from free-living cultures and bacteroids were isolated by the method described above. Auxin levels were subsequently determined using the Salkowski colorimetric assay. Every experiment was performed three times with three technical replicates each time.

### Auxin detection in the periplasmic fraction

To analyse the role of AuxT in phytohormone translocation, periplasmic fractions from *S. fredii* HH103, Δ*auxT* and Δ*auxT*-_p_*auxT* strains were isolated and screened for auxin content. Bacterial cultures, originating from single colonies, were grown in 75 mL of TY medium supplemented with tryptophan (500 mg L^−1^), for 96 hours at 28 °C with continuous shaking.

Periplasmic extraction was performed using a modified osmotic shock and permeabilization protocol (Borrero-de Acuña *et al*., 2015). Briefly, 50 mL of each saturated culture were harvested by centrifugation at 4,000 x g for 20 min at 4°C. Cell pellets were washed with PBS 1X and then gently resuspended, avoiding vortexing to prevent cytoplasmic leakage, in 15 mL of PE buffer [20% (w/v) sucrose and 50 mM Tris (pH 8.0), which was pre-filtered; 1 mg mL □¹ polymyxin B sulfate (Sigma-Aldrich, USA) and protease inhibitor cocktail Mix B (SERVA, Germany)]. The polymyxin and the protease inhibitor cocktail were added immediately prior to use. After overnight incubation with rotation at 4°C, the samples were transferred to 1 mL ultracentrifuge tubes and ultracentrifuged, using a TLA-110 fixed-angle titanium rotor (Optima MAX CP 90NX, Beckman Coulter, USA), at 100,000 x g at 4 °C for 1 h. The resulting supernatant corresponded to the purified periplasmic fraction.

Auxin quantification within these periplasmic extracts was conducted using the Salkowski colorimetric assay as described above. Briefly, 40 µL of periplasmic fractions were mixed with 160 µL of Salkowski reagent and incubated for 20 min in dark. The concentration of indole-3-acetic acid (IAA) and related indolic compounds was determined by measuring absorbance at 535 nm using a plate reader. Every experiment was performed in three biological replicates times with eight technical replicates each time.

To confirm the presence of proteins in the isolated periplasm, 15 µL of each sample were mixed with loading buffer and loaded onto stain-free polyacrylamide gels (Bio-Rad, USA). To rule out cytosolic cross-contamination, a Western blot analysis of the RNA-polymerase was performed comparing the periplasmic and cytosolic fractions of the wild-type HH103 strain. The cytosolic fraction was obtained by resuspending the cell pellet harvested after ultracentrifugation in 1 mL of 50 mM Tris-HCl (pH 8.0). This suspension was transferred to 2 mL tubes containing 0.1 mm glass beads (Scientific Industries, USA) and cells were mechanically disrupted using a FastPrep-24 system (MP Biomedicals, USA). The disruption was carried out by applying 3 cycles at 6.5 m s □¹, with 5-min incubations on ice between cycles to prevent protein degradation.

The lysate was centrifuged at 15,000 x g for 6 min at 4°C. The supernatant was then transferred to a new tube and centrifuged again for 10 min under the same conditions to ensure the complete removal of glass beads and cell debris. The resulting soluble fraction and the periplasmic extract from the strain HH103 were separated by SDS-PAGE and transferred to membranes using the Trans-Blot Turbo Transfer System (Bio-Rad, USA).

Transfer assembly was placed in a Mini Trans-Blot electrophoretic transfer cell (Bio-Rad, USA) and run at 25 V and 1 A for 15 min. The membrane was blocked in 5% (w/v) non-fat milk prepared in TBS-Tween buffer [TBS (10 mM Tris, 150 mM NaCl pH 7,6) containing 0.1% (v/v) Tween 20] for 1 h at room temperature, and was then washed three times for 5 min each with continuous shaking with TBS-Tween buffer at room temperature.

Following this incubation time, the blocking solution was discarded, and the membrane was incubated with anti-*E. coli* RNAPβ (BioLegend, 663903) primary antibody for detection of RNAPβ subunit at a 1:5,000 for 1 hour at room temperature. After three washes with TBS-Tween, HRP-conjugated anti-mouse IgG secondary antibody (Cell Signaling Technology, USA) at 1:5,000 dilution was added for 1 h at room temperature. Protein bands were visualized by applying 250 µL of the chemiluminescent substrate Immobilon Forte HRP (Merck KGaA, Germany). Chemiluminescent signals were detected using the LI-COR Odyssey FC 700/800/Chemi imaging system and analyzed with Image Studio Lite software (LI-COR Biosciences, USA).

### Auxin quantification in the rizosphere

Surface-sterilized seeds of *G. max* var. Williams were germinated and placed on a sterile support grid inside sterilized glass tubes containing Fåhraeus nitrogen-free nutrient solution (Vincent, 1970a). IAA was added to the nutrient solution at a defined concentration (37.5 mg L□¹). Seedlings were then inoculated with 1 mL of bacterial culture at an OD□□□ of 0.6 (approximately 1 × 10□ cells mL□¹). The tubes were incubated in a temperature-controlled growth chamber under the same photoperiod conditions described above for *G. max*. At 3-day intervals, aliquots of the hydroponic solution were collected over a 9-day period, and auxin concentrations were quantified using the Salkowski colorimetric assay. This experiment was performed 3 times with 3 replicates each time.

### Auxin quantification in nodules

To quantify auxin levels within nodules, nodules obtained from nodulation assays of *G. max* var. Williams inoculated with *S. fredii* HH103 Δ*auxT* and Δ*auxT*-_p_*auxT* strains were harvested and homogenized using a mortar and pestle. The resulting homogenate was transferred to 1 mL microcentrifuge tubes and centrifuged at maximum speed for 1 min to pellet plant debris. The supernatants were subsequently subjected to auxin quantification using the Salkowski colorimetric assay. Bacteroids were isolated from nodules using the bEV isolation protocol described above for subsequent quantification of auxin levels. Isolated bacteroids were centrifuged at 12,000 x g for 1 min, and the auxin contained in the resulting supernatants was quantified using the Salkowski colorimetric assay. To quantify auxin levels in bacteroid-derived bEVs, bEVs were isolated as described above, and the resulting bEVs preparations were analysed for auxin content using the Salkowski colorimetric assay.

### Nodulation assays

*S. fredii* HH103 parental strain, Δ*auxT* and Δ*auxT*-_p_*auxT* strains were grown in YM liquid medium. Surface-sterilized seeds of *G. max* var. Williams were germinated and placed in sterilised Leonard jars containing Fårhaeus N-free nutrient solution (Vincent, 1970a). Seeds were then inoculated with 1 mL of bacterial culture at an OD_600_ of 0.6 (10^9^ bacteria mL^-1^). Finally, Leonard jars were placed in a temperature-control chamber with a photoperiod of 16 h at 26 °C in the light and 8 h at 18 °C in the dark, with 70% humidity. Nodulation parameters were evaluated after 6 weeks. The shoots were dried at 70 °C for 48 h and then weighed, and the nodules were harvested from the roots, counted and weighed. Nodulation experiments were performed four times, with five technical replicates for each treatment.

### Statistical analysis

The statistical tests performed in this work are indicated in the figure legends and were done using Prism 8 (GraphPad, USA).

## Results

### The expression of *auxT* transporter and auxin biosynthesis is independent of flavonoid induction in *S. fredii* HH103

Our previous proteomic characterization of bEVs isolated form peribacteroid space established the conceptual framework for this study (Ayala-García *et al*., 2024a). That work revealed an unexpected repertoire of rhizobial proteins potentially involved in host-microbe communication and molecular exchange within this symbiotic niche. Notably, a putative auxin efflux carrier (hereafter AuxT, from Auxin Transporter) was identified as a constituent of the bEVs released by *S. fredii* HH103 bacteroids, suggesting a functional role in phytohormone trafficking.

To perform an initial assessment of the auxin production capacity in HH103 and its potential relationship with the AuxT transporter, we first conducted a series of *in silico* analysis, including genomic-context comparisons, homology searches, orthology assessments, and re-analysis of previously published transcriptomic data (Pérez-Montaño *et al*., 2016). The *auxT* gene (*SFHH103_03783*) is located on the chromosome (**Figure S1A**), whereas the auxin biosynthesis genes *y4wE* (*SFHH103_psfHH103d_257*) and *y4wF* (*SFHH103_psfHH103d_258*) reside on the symbiotic plasmid pD downstream of a canonical *nod* box (NB15; **Figure S1A**) (Vinardell *et al*., 2015). Immediately downstream of *auxT* lies *SFHH103_03784*, annotated as the hypothetical transposase Y4zB, suggesting that *auxT* and *SFHH103_03784* may form a small transcriptional unit. Moreover, the presence of a transposase-associated gene within this region is consistent with a possible acquisition through horizontal gene transfer, potentially contributing to the dissemination of auxin transport functions among rhizobial lineages. Analysis of genomic environment revealed that both genes are flanked by two divergently transcribed genes, *SFHH103_03782* upstream and *ilvE3* (*SFHH103_03785*) downstream, indicating that auxT occupies the first position of an isolated genomic locus (**Figure S1A**).

To further investigate the occurrence of auxin efflux carriers in *S. fredii* HH103, a BLASTP search using AuxT as query identified two additional homologous genes in the genome, *SFHH103_01028* and *SFHH103_01341*, both annotated as members of the AEC family and located in the chromosome. The genomic contexts of these homologs were examined to determine whether they were associated with operons or gene clusters related to auxin metabolism or transport. In neither case did the surrounding genetic organization suggest a direct association with auxin metabolism or transport pathways (**Figure S1B**).

Comparative amino acid sequence analyses revealed different degrees of conservation among the three proteins (**Figure S2A**), supporting their classification within the same transporter family while suggesting potential functional diversification.

To assess the evolutionary conservation of these transporters, orthology and genomic-context analyses were analyzed using GeCoViz (Botas *et al*., 2022). The results showed that all three *S. fredii* HH103 AEC-family transporters are broadly conserved within the *Sinorhizobium*/*Ensifer* clade. Interestingly, *auxT* orthologs were consistently found adjacent to a divergently transcribed gene encoding a BON-domain-containing protein (**Figure S2B**). In *S. fredii* HH103, this corresponds to *SFHH103_03782*, the gene immediately upstream of *auxT*. BON-domain proteins have been associated with membrane stress responses and cell-envelope remodeling processes (Yeats & Baterman, 2003). Given that AuxT was identified as a component of *S. fredii* HH103-derived bEVs, the conserved genomic association between *auxT* and the BON-domain gene might reflect a functional link to membrane remodeling mechanisms involved in bEVs biogenesis or protein incorporation into vesicles, although this possibility remains to be experimentally addressed.

Re-analysis using published RNA-seq data (Pérez-Montaño *et al*., 2016) revealed that *auxT* expression under free-living conditions is independent of the *nod* inducing flavonoid genistein, remaining its transcript levels moderate (rank ∼4,600 of the total 7,014 genes) regardless of flavonoid presence. Conversely, *y4wE* exhibited strong flavonoid-dependent induction (24.7-fold, rising from rank 4,666 to 185), while *y4wF* showed constitutively very high expression (rank 255 without flavonoids, 112 with flavonoids, 2-fold) (**Figure 1A, Table S4**). These data demonstrate that while *y4wE* is induced by NodD1 via NB15, both *auxT* and specially *y4wF* are constitutively expressed independently of flavonoid presence.

**Figure 1.**
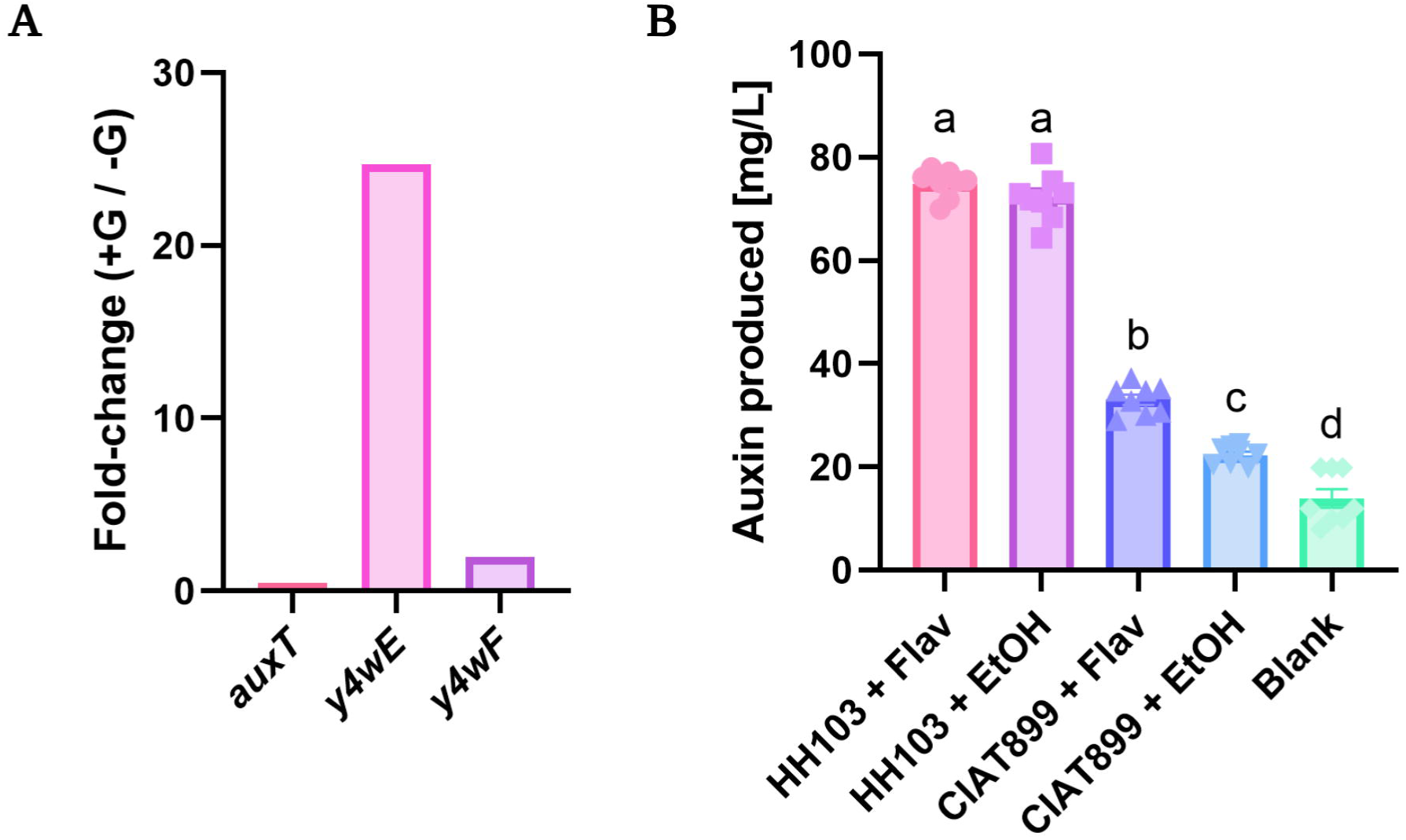
**A)** Relative expression of *auxT* and the auxin biosynthesis genes *y4wE* and *y4wF* in response to flavonoid treatment. Data were obtained from previous RNA-seq studies performed by Pérez-Montaño *et al*., (2016). Transcript levels are expressed as fold change relative to cultures grown in the absence of flavonoids. Differentially expressed genes were defined as those genes with a fold-change lower or higher than − 3 or 3, respectively, with a *p*-value lower to 0.05 (3 biological replicates). **B)** Auxin production by *S. fredii* HH103 and *R. tropici* CIAT 899 in the presence and absence of flavonoids. The blank corresponds to TY-tryptophan medium without bacterial inoculation. Values represent the mean ± SD of 3 independent biological replicates with at least 5 technical replicates. Bars labeled with the same letter are not significantly different according to one-way ANOVA (α < 0.0001). Flav: Flavonoid; EtOH: Ethanol.

Next, we quantified auxin production in *S. fredii* HH103 using the Salkowski colorimetric assay under both inducing (flavonoid presence) and non-inducing conditions (flavonoid absence). Notably, similar to *S. fredii* NGR234 (Theunis *et al*., 2004), this strain exhibited significant constitutive auxin production independently of the presence of flavonoids. These values exceeded those of the control strain, the inducible producer *R. tropici* CIAT 899, under both inducing and non-inducing conditions (**Figure 1B**).

### AuxT adopts a PIN-like architecture with structural homology to plant auxin transporters

Next, we set out to elucidate the *in silico* basis of the three-dimensional structure of AuxT. We employed AlphaFold3 to predict the three-dimensional structure of AuxT. Notably, the first 87 amino acids of the originally annotated AuxT sequence are absent in all orthologs of the *Sinorhizobium* family and lack defined secondary structure in our models, suggesting they correspond to an annotation error in the published genome as aforementioned (Vinardell *et al*., 2015). Consequently, all structural analyses were performed using the corrected 314-amino acid sequence. The model revealed a compact, multi-pass membrane protein organized into ten transmembrane α-helices arranged in two pseudosymmetrical bundles (**Figure 2AB**), a topology characteristic of the AEC family (Ung *et al*., 2022). This topology was supported by DeepTMHMM v1.0, which predicted ten transmembrane helices connected by alternating cytoplasmic and periplasmic loops. This protein was predicted to possess a cytoplasmic orientation for both its N- and C-termini. SignalP v6.0 analysis detected no secretory signal peptide, ruling out SecYEG- or Tat-dependent export and supporting localization as an integral membrane protein. PSORTb v3.0 assigned AuxT to the cytoplasmic membrane with maximum confidence, further validating its predicted subcellular location.

**Figure 2.**
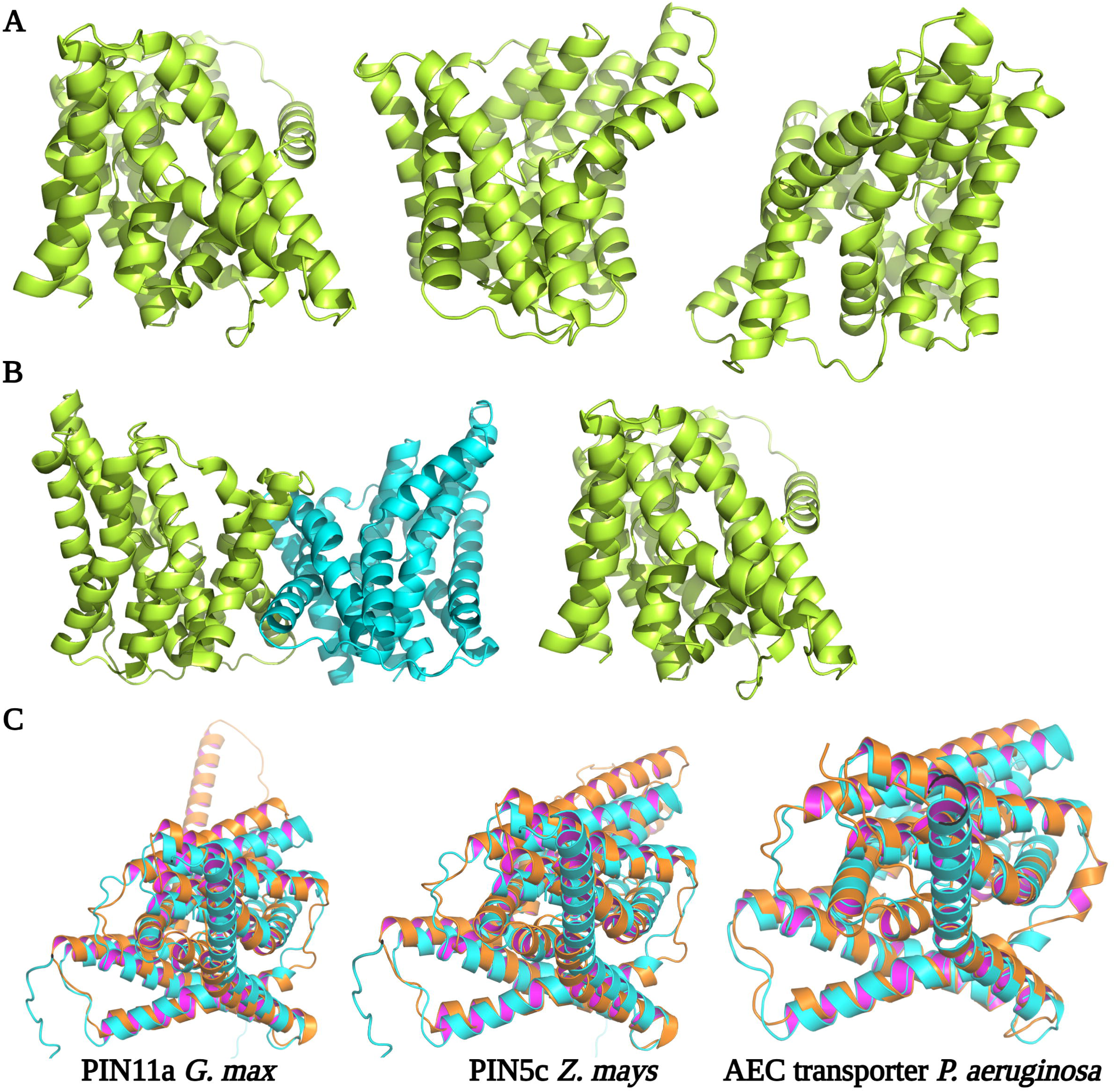
**A)** AlphaFold3 cartoon model of the AuxT monomer (green) shown in front, right-side, and left-side views, respectively. **B)** AlphaFold3 cartoon model of the predicted AuxT dimer, consisting of monomer A (green) and monomer B (cyan), shown in front and side views, respectively. The dimeric interface was validated by high computational confidence scores (ipTM = 0.85; average PAE < 5 Å). **C)** Structural alignment of AuxT (cyan) with auxin transporters (orange) from *G. max* PIN11a, *Z. mays* PIN5c, and a putative auxin efflux carrier (AEC) transporter from *P. aeruginosa*. Note that the inwards directed side of helices is colored in pink. Structural superpositions yielded RMSD values of 3.52 Å over 1,416 aligned atoms, 3.86 Å over 1,494 aligned atoms, and 1.75 Å over 1,359 aligned atoms, respectively. The near-identical RMSD of 1.75 Å with the *P. aeruginosa* AEC transporter highlights AuxT’s structural conservation within its bacterial family, while the higher RMSD values (>3.5 Å) with plant PIN transporters indicate a conserved transmembrane topology with variations in rigid-body orientation and loop regions.

To identify structural homologs, we performed a structure-based search using Foldseek against the AlphaFold Database. Remarkably, despite negligible sequence identity, the top hits were not bacterial proteins but plant PIN-family auxin transporters. These included PIN11a from the natural host *G. max* (ID Uniprot: I1K2H7; E-value 2.82e-10), PIN5c from *Z. mays* (ID Uniprot: B4FSJ4; E-value 2.30e-11) and PIN5 from *A. thaliana* (ID Uniprot: Q9FFD0; E-value 5.54e-10). Even more, AuxT exhibited very high structural similarity to an AEC family transporter from *Pseudomonas aeruginosa* (ID UniProt: Q9HTA4; E-value 3.35e-12), a protein from a plant-associated bacterium (**Table S5**).

To further evaluate the structural relationships between AuxT and the proteins identified by Foldseek, we performed pairwise structural alignments using the AlphaFold3 model of AuxT and representative homologs from *G. max*, *Z. mays* and *P. aeruginosa*. The AlphaFold3-predicted dimeric model of AuxT was additionally validated by high interface prediction scores, with an interface predicted TM-score (ipTM) of 0.85 and low Predicted Alignment Error (PAE) values across the dimer interface (average PAE < 5 Å), providing strong computational support for the biological relevance of the dimeric assembly. Structural similarity was assessed using Root Mean Square Deviation (RMSD) values measured in Ångströms (Å). RMSD values between 2 and 4 Å over multi-pass transmembrane proteins indicate a recognized topological fold while also reflecting rigid-body orientation or loop variations, whereas values below 2 Å indicate near-identical structures. RMSD values were interpreted together with the number of aligned atoms, as larger alignments provide greater statistical confidence.

Structural alignment with *G. max* PIN11a yielded an RMSD of 3.52 Å over 1,416 aligned atoms (**Figure 2C**), evidencing conserved topological features of the transmembrane core despite overall sequence divergence. A similar result was obtained for *Z. mays* PIN5c, which aligned with an RMSD of 3.86 Å over 1,494 atoms, further supporting structural conservation among PIN-family transporters. In contrast, alignment with the *P. aeruginosa* AEC transporter produced an RMSD of 1.75 Å over 1,359 aligned atoms (**Figure 2C**), indicating near-identical folding of the transmembrane core and demonstrating that AuxT remains structurally tight within its bacterial family context while mimicking plant transporters topologically. In all cases, the conserved transmembrane bundle and the predicted inward-facing cavity were maintained.

### Molecular docking reveals auxin binding to AuxT

To bioinformatically test auxin-binding capacity and capture ligand-induced conformational changes, we generated ligand-bound structural models of monomeric and dimeric AuxT for a panel of auxin-family ligands (IAA, 4-Cl-IAA, IBA, and PAA) using the RFAA (RoseTTAFold All-Atom) co-folding platform. Unlike classical rigid-backbone docking programs (e.g., AutoDock Vina), RFAA performs full-atom co-modeling of the protein and small molecule, allowing both the protein backbone and side-chains to undergo structural rearrangements upon ligand binding. The tetrameric form was excluded, as eukaryotic structural homologs do not exhibit biological activity in this oligomeric state (Ung *et al*., 2022). To quantify the extent of these conformational changes, we calculated the Cα RMSD between the ligand-free (apo) and ligand-bound (holo) structural models.

In the monomeric form, IAA bound within the central inward-facing cavity formed by transmembrane helices 3, 4, 6, and 7, with key interactions involving residues W192, Y194, and F361. However, ligand binding induced only minimal Cα backbone rearrangements in the monomer, with RMSD values below 1.36 Å across all tested ligands (**Figure 3A**). This lack of substantial structural change indicates that the monomer adopts a rigid, resting-state conformation that lacks the conformational flexibility required for active transport. In stark contrast, the AuxT dimer underwent profound and ligand-specific conformational changes upon ligand binding. IAA and 4-Cl-IAA binding induced large-scale structural rearrangements with Cα RMSD values of 14.40 Å and 13.38 Å, respectively, with the ligand positioned at the dimer interface (**Figure 3B**). PAA induced the most dramatic shift, yielding a Cα RMSD of 16.56 Å, while IBA led to a more modest "closed" conformation with an RMSD of 5.75 Å. The marked contrast between the minimal monomeric RMSDs (< 1.36 Å) and the extensive dimeric RMSDs (up to 16.56 Å) highlights that dimerization is essential for the transporter’s conformational dynamics and ligand responsiveness. These large-scale movements are consistent with the elevator-type transport mechanism described for plant PIN8 (Ung *et al*., 2023).

**Figure 3.**
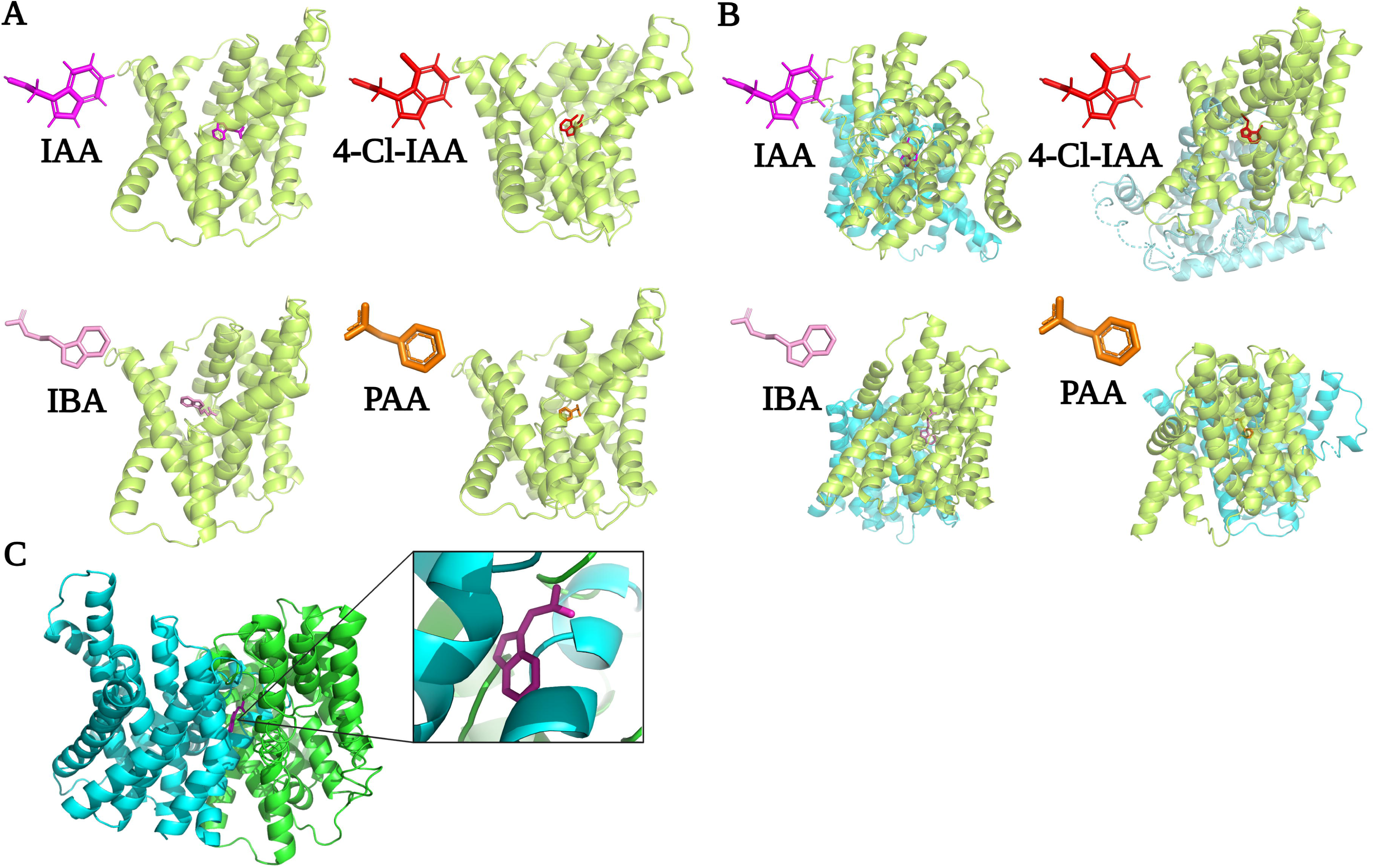
**A)** Docking of the AuxT monomer (green) with IAA (magenta), 4-Cl-IAA (red), IBA (pink) and PAA (orange). Predicted docking poses yielded low RMSD values: 1.36 Å, 1.21 Å, 1.26 Å and 1.34 Å, respectively. **B)** Docking of the predicted AuxT dimer, composed of subunit A (green) and subunit B (cyan), with the same auxin-related compounds. RMSD values were 14.40 Å for IAA, 13.38 Å for 4-Cl-IAA, 5.75 Å for IBA, and 16.56 Å for PAA. **C)** Structural model of the AuxT dimer (cyan and green) interacting with a generic auxin molecule (magenta) represented by an aromatic ring linked to a carboxyl-containing side chain, illustrating the proposed auxin-binding cavity.

To complement these simulations, an AlphaFold model of the AuxT dimer with a generic auxin molecule showed a compact conformation where the ligand is localized between the two subunits, interacting with both (**Figure 3C**). This predicted geometry is compatible with an alternating-access transport mechanism described elsewhere (Ung *et al*., 2022). These results identify the dimer as the conformationally dynamic, and likely functional, state of AuxT. The ligand-dependent nature of conformational changes suggests that AuxT can distinguish between different auxin species, with potential implications for substrate specificity *in vivo*.

### The rhizobial AuxT transporter is highly enriched in bacterial extracellular vesicles

After demonstrating constant auxin production, constitutive *auxT* expression, and *in silico* evidence for AuxT-mediated auxin transport in *S. fredii* HH103, we investigated whether AuxT is loaded into bEVs at biologically relevant levels to clarify the potential role of AuxT in vesicle-mediated auxin transport. To this end, we first quantified bEV yields and subsequently confirmed the presence of AuxT within the vesicular cargo of *S. fredii* HH103. To perform these experiments, we generated an *auxT* non-polar deletion mutant (Δ*auxT*) by double homologous recombination. For complementation, we introduced the pSEVA221::P*auxT*-*auxT*::*streptagII* plasmid into the mutant, yielding Δ*auxT-*_p_*auxT*, which expresses C-terminal StrepII-tagged *auxT* governed by its native promoter. It is worth noting that this plasmid is maintained at approximately 4-7 copies in *S. fredii* (Silva-Rocha *et al*., 2013; Thomas *et al*., 1984), which result in moderately increased global expression of *auxT* despite being under the control of its native promoter.

Field emission scanning electron microscopy revealed abundant cell surface blebbing consistent with bEV biogenesis (**Figure S3A**). Purified bEVs from free-living cultures and bacteroids appeared as intact spherical structures (40-200 nm) by transmission electron microscopy (**Figure S3BC**). Deeper characterization by Nanoparticle Tracking Analysis (NTA) revealed that bEV production differs between free-living and symbiotic conditions. Free-living cultures produced 1.85 × 10¹¹ ± 2.95 × 10¹□ particles/mL in wild-type HH103, 1.54 × 10^11^ ± 1.9 × 10^10^ particles/mL in Δ*auxT* and 1.6 × 10^10^ ± 1.2 × 10^9^ particles/mL in Δ*auxT-*_p_*auxT* (**Figure 4A**), while bacteroid-derived bEVs yielded 4.96 × 10^10^ ± 2.56 × 10^9^ particles/mL in wild-type HH103, 2.69 × 10^10^ ± 1.25 × 10^9^ particles/mL in Δ*auxT* and 1.68 × 10^10^ ± 7.63 × 10^8^ particles/mL in Δ*auxT-*_p_*auxT* (**Figure 4B**). The most frequent particle size in free-living bEVs was 81.6 ± 9.7 nm for wild-type HH103, 63.3 ± 11 nm for Δ*auxT* and 172.5 ± 13.3 nm for Δ*auxT-*_p_*auxT* (**Figure 4C**); whereas bacteroid-derived bEVs were larger, with 163.2 ± 6.9 nm for wild-type HH103, 136.6 ± 6.6 nm for Δ*auxT* and 150.5 ± 10.2 nm for Δ*auxT-*_p_*auxT* (**Figure 4D**), in alignment with previous observations (Ayala-García *et al*., 2024a). When statistical analysis was performed, no significant differences in bEV concentration were observed between wild-type, Δ*auxT*, or Δ*auxT-*_p_*auxT* strains under free-living conditions nor on those derived from the bacteroid (**Table 1**).

**Figure 4.**
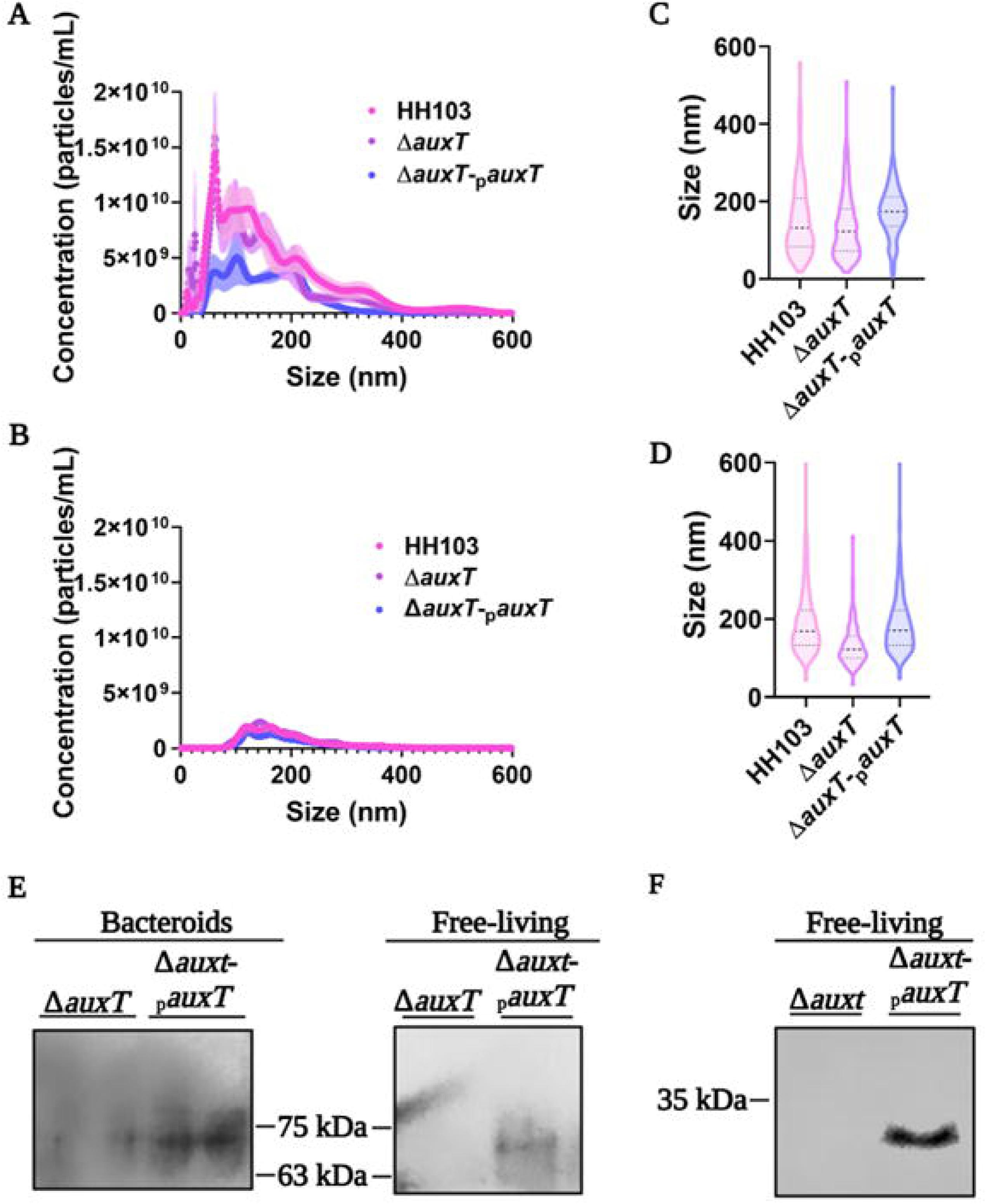
**(A-B)** Particle concentration (particle/mL) profiles obtained by NTA of bEVs isolated from free-living cultures (A) and soybean bacteroids (B), showing particle concentration as a function of vesicle size. The shaded regions around the curves represent the variation among biological replicates **(C-D)** Violin plots summarizing vesicle size distributions obtained by NTA for bEVs derived form free-living cultures (C) and bacteroids (D). **(E-F)** Detection of AuxT in bEVs by western blot analysis using the Δ*auxT* mutant and the complemented strain Δ*auxT*-_p_*auxT*. (E) Detection of the predicted AuxT dimer in bEVs isolated both from free-living cultures and bacteroids. (F) Detection of the AuxT monomer in bEVs isolated from free-living cultures. Relative molecular weight is indicated in kDa.

**Table 1.**
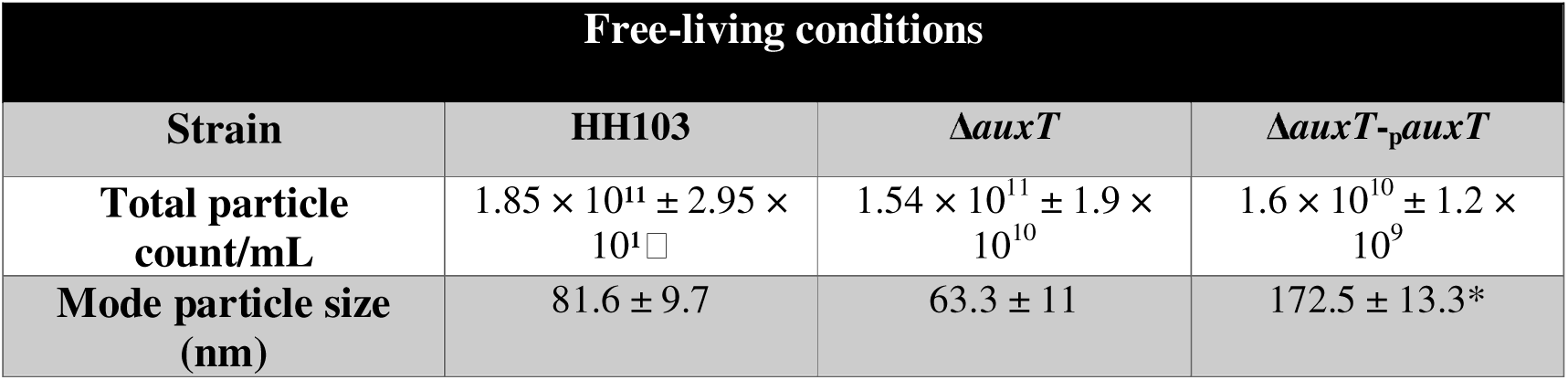
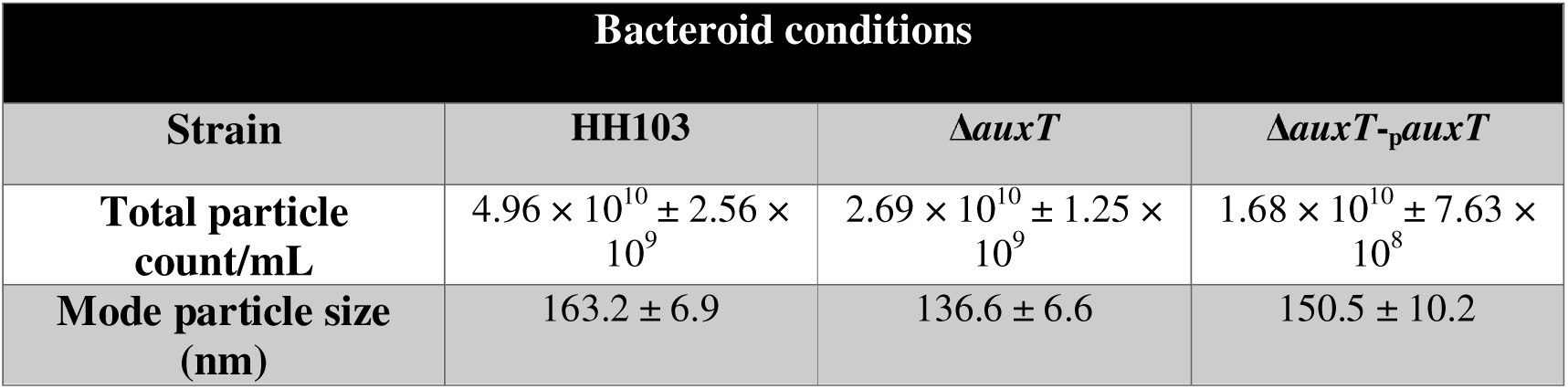
Characterization by NTA of bEVs isolated from the parental strain of *S. fredii* HH103, the Δ*auxT* mutant and the complemented strain Δ*auxT-*_p_*auxT* under free-living and bacteroid conditions. The table summarizes the total particle count and the mode particle size of each vesicle preparation. Statistical comparisons were performed using one-way ANOVA. Significance levels regarding to the wild-type strain (HH103) are indicated as * (α=0,05).

Despite finding no statistically significant differences in size and concentration in bacteroid conditions, the Δ*auxT-*_p_*auxT* strain exhibits a distinct size-distribution profile characterized by a shift toward larger vesicle sizes under free-living conditions (**Table 1**). In fact, the NTA profile reveals a pronounced distribution toward 200 nm, indicating an increased production of a larger vesicle subpopulation compared to the wild-type and Δ*auxT* strains. This effect may be explained by the higher abundance of AuxT transporters embedded in the inner membrane, which likely enhances auxin translocation into the periplasm. The resulting accumulation of auxins in the periplasm could promote membrane blebbing and stimulate the formation of larger bEVs with an increased capacity to transport auxins, which are poorly soluble and possess acidic, potentially harmful properties (Jemiola- Rzemińska *et al*., 2015). Indeed, hydrophobic compounds and other chemically harsh molecules that disrupt cell homeostasis and accumulate in the periplasm have previously been reported to stimulate OMV formation, as vesiculation serves as a mechanism to facilitate their secretion from the cell (Toyofuku *et al*., 2023).

Subsequently, we performed immunoblot analysis employing 1 × 10^9^ bEVs obtain from each strain and culture condition. In the bEVs from the Δ*auxT-*_p_*auxT* strain, obtained under free-living conditions and from fully differentiated bacteroids, a specific immunoreactive band at ∼70 kDa was immunodetected, which was absent in bEVs from the Δ*auxT* mutant (**Figure 4E**). Although gene annotation predicts a monomeric protein of ∼35 kDa, comparative analysis of AuxT across rhizobial strains (see above) indicates a misannotation in *S. fredii*, with the functional protein likely initiating at amino acid 88 and thus having a corrected molecular weight of ∼33 kDa. The predominance of a ∼70 kDa band under standard denaturing conditions, rather than the expected ∼33 kDa monomer, suggests that AuxT forms a highly stable dimer. This stability may reflect its transmembrane nature, rich in α-helices that promote SDS binding and anomalous migration (Rath *et al*., 2009), as well as the presence of a conserved cysteine residue per monomer, potentially enabling interchain disulfide bond formation (Bavoil *et al*., 1984; Fukuda *et al*., 2001). Under more stringent denaturing conditions consisting of prolonged heating, a ∼33 kDa band corresponding to the monomer became detectable (**Figure 4F**), confirming the revised molecular weight.

Following, we visually confirmed AuxT occurrence within legume nodules by confocal microscopy. To this end, we constructed an N-terminal *m*Cherry-AuxT fusion under the control of its native promoter. Confocal microscopy of *Lotus burttii* nodules harbouring this construct revealed a soft red fluorescence localized to the infection zone, overlapping with bacteroid-occupied cells. Control nodules transformed with the empty vector exhibited only background autofluorescence (**Figure S5**). This microscopic inspection confirmed that AuxT is substantially produced within nodules during symbiosis. Collectively, these results demonstrate that AuxT is a *bona fide* rhizobial bEV cargo under free-living conditions and, crucially, *in planta*, where it may play a substantial role in phytohormone trafficking within the peribacteroid space i.e. from bacterial cells to the plant host cell, facilitating auxin delivery across the symbiotic interface.

### AuxT-mediated auxin loading into bEVs underpins phytohormone trafficking *in planta* and in free-living conditions

Auxins are negatively charged under physiological conditions. This, coupled with their bulky aromatic structure, makes the presence of active transporters necessary for them to cross membranes (Becker *et al*., 2020). We therefore hypothesized that the inner membrane-embedded transporter AuxT functions to translocate auxins from the cytoplasm into the periplasm, where the phytohormone could subsequently be incorporated into the bEVs, mainly B-OMVs which are known to carry periplasmic and outer membrane content (Toyofuku *et al*., 2023). Although, auxin incorporation into B-OIMVs represent a possibility as well. To first determine whether AuxT directly contributes to auxin loading into bEVs, we purified vesicles from both free-living cultures and bacteroids of the Δ*auxT* and Δ*auxT*-_p_*auxT* strains. Consistent with our hypothesis, bEVs from the complemented strain contained significantly higher auxin levels than those derived from the mutant under both conditions (**Figure 5A and 5B**), confirming that AuxT facilitates auxin encapsulation into these vesicles. Nevertheless, the presence of residual auxin in bEVs from the Δ*auxT* background suggests the existence of AuxT-independent transport mechanisms. As aforementioned, two additional *auxT* homologs are present in the genome of *S. fredii* HH103. However, their encoded proteins were not detected in the bEV proteome in our previous study, suggesting that they are not sorted into the selected cargo of bEV (Ayala-García *et al*., 2024a).

**Figure 5.**
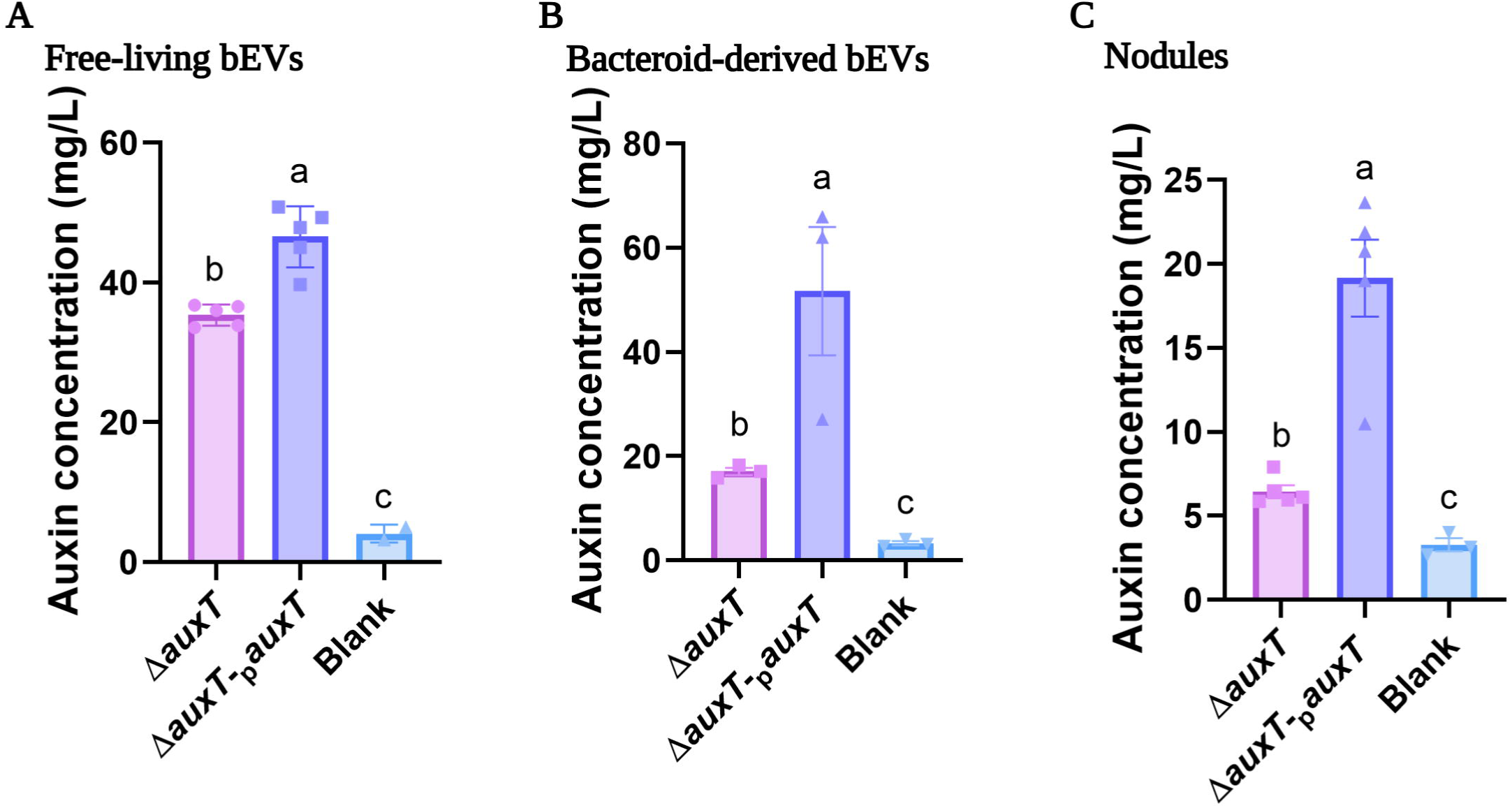
**(A-B)** Auxin concentration in bEVs isolated from free-living cultures **(A)** and from bacteroids **(B)**. bEVs were isolated from the Δ*auxT* mutant and the complemented strain Δ*auxT*-_p_*auxT*. **(C)** Auxin concentration in soybean nodules inoculated with Δ*auxT* and Δ*auxT*-_p_*auxT* strains. Saline solution was used as a blank control. Values represent the mean ± SD of 3 independent biological replicates with at least 5 technical replicates. Bars labeled with the same letter are not significantly different according to one-way ANOVA (α < 0.0001).

To further assess whether AuxT functions as an inner membrane exporter required for auxin translocation into the periplasm prior to bEV loading, we quantified auxin accumulation in the periplasmic fraction of the wild-type, Δ*auxT* mutant, and the complemented strain Δ*auxT*-_p_*auxT* using the Salkowski colorimetric assay. To confirm the absence of cytosolic cross-contamination in the periplasmic extracts, we performed a western blot using RNA polymerase as a cytosolic marker in both the periplasmic fractions and the corresponding cell pellets of the wild type strain. The periplasmic fractions contained abundant proteins in the SDS-PAGE gel but showed no detectable RNA polymerase signal by immunodetection, in contrast to the pellet control, where the marker was clearly detected, indicating the absence of intercompartmental contamination (**Figure S4A**). As expected, the wildtype presented the highest accumulation of auxins in the periplasm (**Figure S4B**). Interestingly, the Δ*auxT* mutant still accumulated substantial amounts of auxin in the periplasm, indicating that AuxT homologs likely contribute to auxin translocation across the inner membrane (**Figure S4B**). Strikingly, the Δ*auxT* mutant exhibited significantly higher auxin accumulation in the periplasm than the complemented strain showed (**Figure S4B**). A possible explanation is that transporter overexpression in the complemented strain dramatically increases auxin translocation into the periplasmic space. This homeostatic disruption could induce the release of auxin-loaded bEVs to mitigate the stress induced by phytohormone accumulation, reducing the total concentration of these molecules within the periplasmic space (Lima *et al*., 2022).

Altogether, these observations support a model in which AuxT actively mobilizes auxin across the inner membrane into the periplasm, where local auxin accumulation stimulates bEV formation and enables its subsequent packaging for export and delivery to the host plant.

Once AuxT-dependent auxin loading into bEVs was established under both free-living and bacteroid conditions, we assessed whether this translated to a higher concentration of the phytohormone within the whole nodule by quantifying *in planta* auxin levels. Remarkably, nodules from plants inoculated with the Δ*auxT*-_p_*auxT* accumulated significantly higher auxin levels than those from the Δ*auxT* mutant (**Figure 5C**). In conjunction with the previous immunoblotting assays and confocal microscopy studies, these results confirm that AuxT is present in the symbiosomes of legume nodules induced by *S. fredii* HH103 and functions to mobilize auxins across the inner membrane, enabling their subsequent loading into bEVs secreted at the plant-microbe interface.

### AuxT is required for an optimal symbiotic performance

Building on evidence from both *in silico* and *in vivo* analyses identifying AuxT as a functional auxin transporter within bEV in the peribacteroid interface, we next addressed whether this phytohormone transport pathway contributes to the symbiotic efficiency of *S. fredii* HH103. To assess the physiological relevance of AuxT, we performed nodulation assays using *G. max* cv. Williams. The Δ*auxT* strain exhibited significant symbiotic defects compared to the wild-type and complemented strains. Specifically, plants inoculated with the Δ*auxT* mutant strain showed significant reduced shoot dry mass (1.3–1.4-fold decrease; **Figure 6A**), nodule number (1.5-fold decrease; **Figure 6B**), and total nodule mass (1.4–1.5-fold decrease; **Figure 6C**) compared with those inoculated with the wild-type and complemented strains. These parameters were fully restored to wild-type levels in the Δ*auxT*-_p_*auxT* complementary strain, confirming that the phenotype is attributable to the loss of *auxT*. Consistently, visual inspection revealed smaller plants with fewer nodules in the mutant treatment (**Figure 6D**). Altogether, these findings demonstrate that AuxT-mediated auxin packaging within bEVs derived from the *S. fredii*-derived bacteroid contributes to the symbiotic efficiency of *S. fredii* HH103, significantly impacting overall plant growth promotion.

**Figure 6.**
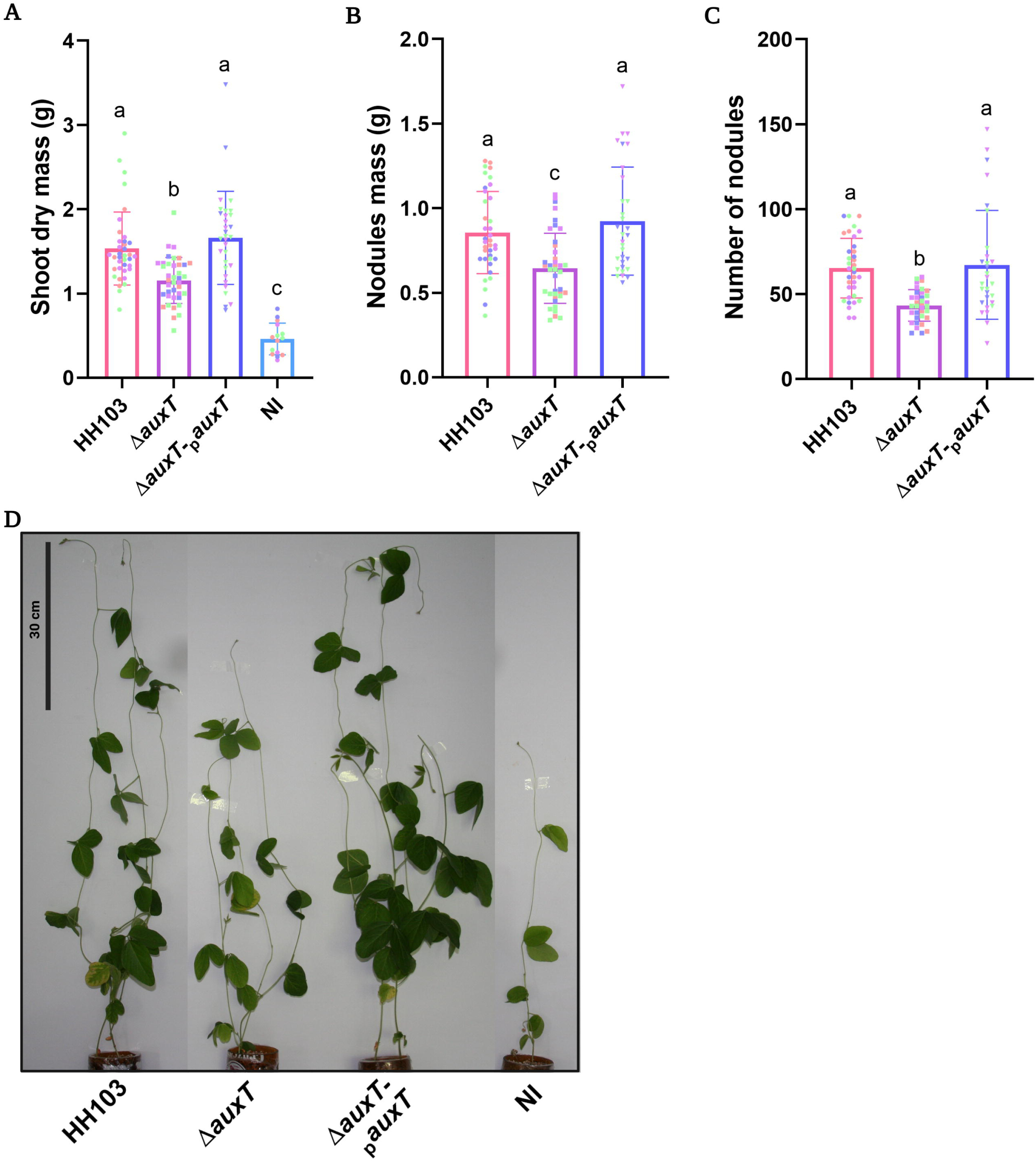
Soybean responses to inoculation with the parental strain *S. fredii* HH103, the Δ*auxT* mutant or the complemented strain Δ*auxT*-_p_*auxT*. **(A)** Shoot dry mass. **(B)** Nodule mass. **(C)** Number of nodules per plant. Data are the mean ± standard error of the mean per plant. Individual points represent values from single plants, and colors indicate independent nodulation experiments. Values represent the mean ± SD of 3 independent biological replicates with at least 5 technical replicates. Bars labeled with the same letter are not significantly different according to one-way ANOVA (α < 0.0001). **(D)** Shoot development of soybean plants inoculated with the different strains. NI: non-inoculated.

## Discussion

The intricate molecular dialogue underpinning the rhizobium-legume symbiosis is crucial for establishing efficient nitrogen-fixing interactions. Our study significantly expands this paradigm by demonstrating a novel role for rhizobial EVs as delivery vehicles for auxins, a process mediated by a PIN-like auxin transporter that actively modulates phytohormone dynamics at the symbiotic interface. To our knowledge, AuxT represents the first described PIN-like auxin transporter discovered in bacteria and packed into bEVs. While rhizobial auxin biosynthesis has been well-documented for over three decades (Costacurta & Vanderleyden, 1995; Patten & Glick, 1996), the precise machinery governing bacterial auxin export during the symbiotic process had remained largely unexplored. The discovery of AuxT successfully fills this physiological gap, revealing that rhizobia possess dedicated membrane proteins for directional bEVs-based auxin transport in symbiosis that are functionally analogous to the plant PIN systems that control endogenous auxin distribution (Ung *et al*., 2023).

In our pioneering study led by Ayala-García *et al*. (2024a), bacteroid-produced bEVs were isolated and characterized for the first time. This work demonstrates that these bEVs contain a broad protein repertoire potentially involved in symbiosis, highlighting novel participants in the rhizobia-plant molecular dialogue. Given that the peribacteroid space is an unhindered interdomain interface between the plant cell and the bacteroid, at least part of the bacteroid bEV cargo is likely destined for host-plant communication. In this context, we focused on AuxT, a structural homologue of the AtPIN8-like auxin transporter whose presence in bEVs was experimentally observed. Bioinformatic analyses indicate that AuxT belongs to the AEC superfamily, a group of bacterial membrane transporters associated with aromatic compound transport. Interestingly, while sequence similarity aligns AuxT with conserved bacterial rhizobial transporters, structural comparisons reveal high homology to plant PIN-family auxin transporters. This structural divergence suggests that AuxT retains a functional architecture for organic small-molecule transport similar to its plant counterparts. This interpretation aligns with the recent proposal by Vosolsobě *et al*. (2024), who suggested that the AEC superfamily emerged early in bacteria as a set of transporters involved in organic acid trafficking, and that certain members subsequently evolved into the PIN and PILS auxin transporter families in plants. Within this framework, it is noteworthy that specific rhizobia have evolved the capacity to transport auxins, likely as a functional adaptation to modulate host-plant interactions and promote symbiosis. The prediction of multiple transmembrane domains, coupled with its localization to the cytoplasmic membrane, strongly supports its proposed transport function (**Figure 2**). Moreover, the predicted multi-pass architecture of AuxT is characteristic of proteins mediating small-molecule translocation, reinforcing the hypothesis that this protein participates in metabolite exchange between the bacterial cell and its immediate environment (Pizzagalli *et al*., 2021).

Genomic context and transcriptional regulation analyses suggest that *auxT* is not part of the classical auxin biosynthesis module described for *S. fredii* HH103. While the genes involved in auxin production, *y4wE* and *y4wF*, are located on the symbiotic plasmid pD and are located downstream a *nod* box, *auxT* is situated on the chromosome, physically separated from this biosynthetic cluster (**Figure S1**) (Vinardell *et al*., 2015). This genomic uncoupling indicates that its function is likely linked to transport or redistribution processes rather than directly to flavonoid-induced auxin synthesis. The observed expression profiles support this hypothesis; *y4wE* (but not *y4wF*) is strongly induced by flavonoids (**Figure 1**), consistent with its NodD-dependent regulation during early symbiotic stages, whereas *auxT* exhibits a moderate but constitutive expression level independent of these inducing compounds (Pérez-Montaño *et al*., 2016). Furthermore, the presence of a hypothetical transposase gene in its immediate genomic surroundings suggests that the *auxT* locus may have been acquired via horizontal gene transfer, an idea reinforced by *in silico* sequence comparisons showing *auxT* homologs in phylogenetically distant and non-symbiotic bacterial species (not shown), which would explain its relatively isolated genomic organization relative to other auxin metabolism genes of *S. fredii* HH103 (Schmitz & Querques, 2024). Conversely, evolutionary conservation and genomic context analyses allowed us to explore the distribution of *auxT* and its orthologs across representative species of the *Rhizobiaceae* family (**Figure S2**). This approach revealed that this gene is widely conserved specifically within the *Sinorhizobium/Ensifer* lineage, suggesting it may perform relevant, conserved physiological functions in these rhizobia.

Structural analysis of AuxT provides additional evidence supporting its involvement in auxin transport. First, three-dimensional models predicted by artificial intelligence and machine learning reveal a typical multi-pass membrane transporter architecture, reinforcing the localization predictions. Furthermore, a remarkable structural similarity was observed between AuxT and auxin transporters characterized in *Glycine max* and *Zea mays* (**Figure 2**). Despite sequence divergence, structural alignment revealed significant conservation, suggesting a preservation of similar functional mechanisms. This aligns with previous studies highlighting that structural homology can be maintained over extensive evolutionary periods despite sequence divergence (Hamamsy *et al*., 2024). Therefore, the architecture of AuxT is compatible with a transport mechanism similar to that characterized for plant PIN-family transporters. PIN transporters, the primary mediators of polar auxin transport in plants, function as homodimers organized into two functional domains: a scaffold domain that stabilizes the dimer interface, and a transporter domain responsible for substrate translocation. Auxin transport occurs via an elevator-type mechanism, where conformational changes within the transporter domain are driven by the negative charge of the auxin molecule itself, operating in coordination with electrochemical gradients (Ung *et al*., 2023). Molecular docking analyses strengthened this possibility by showing that various molecules of the auxin family can interact with AuxT within the predicted structural models (**Figure 3**). Notably, these simulations suggest that the dimeric conformation of the protein undergoes significantly more pronounced conformational changes upon ligand binding compared to the monomeric form (**Figure 3**). This behavior suggests that dimerization may be necessary for the functional activity of the transporter, mirroring plant PIN transporters, which arrange into homodimers (Ung *et al*., 2023). In fact, the difficulty obtaining the monomeric form by western blot in our assays supports this view (**Figure 4**).

Characterization of the size and concentration of bEVs isolated from the parental, *auxT* mutant, and complemented strains revealed no noticeable differences under bacteroid conditions. However, distinct variations emerged in the size distribution of bEVs produced under free-living conditions (**Figure 4**). Specifically, bEVs from the complemented strain exhibited a considerably larger mode particle size. This effect could be linked to an increased abundance of AuxT transporters in the inner membrane; the complemented strain carries a plasmid maintained at four to seven copies *per* cell in *S. fredii* HH103, potentially leading to higher *auxT* expression levels than in the wild-type strain. Consequently, a higher density of AuxT could enhance auxin accumulation within the periplasmic space, thereby promoting vesiculation and the release of larger bEV amounts. The accumulation of auxins, which are negatively charged at physiological pH, is known to destabilize membranes and alter electrochemical gradients, among other biophysical effects (Chimerel *et al*., 2012; Labeeuw *et al*., 2016). Consequently, their rapid bEV-mediated translocation from the periplasm to the extracellular milieu would help mitigate the physiological stress caused by this agent. Western blot analyses confirmed the presence of AuxT in both free-living and bacteroid-derived bEVs, demonstrating that AuxT is indeed part of the vesicular cargo in *S. fredii* HH103 (**Figure 4**). Notably, the protein was predominantly detected in its dimeric form, suggesting that this conformation may be highly stable even under standard denaturing conditions (**Figure 4**). Furthermore, confocal microscopy confirmed the presence of AuxT within the nodule, which indicate that it is actively expressed during this symbiotic stage (**Figure S5**).

Results from auxin production assays confirmed that *S. fredii* HH103 possesses a high capacity for auxin synthesis, reaching levels superior to those detected in *Rhizobium tropici* CIAT899 (**Figure 1**), a species well-known for its flavonoid-induced auxin production (Figueiredo *et al*., 2008). Despite the upregulation of one of the two auxin biosynthesis genes by inducing flavonoids, phytohormone production in *S. fredii* HH103 appears independent of these signals, given that total auxin levels remain unaltered but high in both the presence and absence of genistein. This aligns with previous transcriptomic data demonstrating that the *y4wF* gene is highly and constitutively expressed independently of the presence of these symbiotic molecules (Pérez-Montaño *et al*., 2016). In this scenario, a transport system such as AuxT could be particularly relevant for modulating auxin distribution within the bacteria-plant interaction zone. Auxin quantification in bEVs supports this possibility, as vesicles isolated from the complemented strain exhibit a significantly higher auxin concentration than those from the *auxT* mutant (**Figure 5**). This finding suggests that AuxT participates in loading or transporting these molecules into bEVs. Nevertheless, the detection of auxins in the mutant strain bEVs indicates that this mechanism is not exclusive, and alternative pathways likely contribute to the incorporation of indolic compounds into these structures. In this regard, proteins encoded by the *auxT* homologs identified in the *S. fredii* HH103 genome could represent alternative mechanisms facilitating vesicular auxin loading. In line with these findings, nodule analyses strongly reinforce this interpretation. Thus, nodules from plants inoculated with the complemented strain, as well as the bEVs isolated from their bacteroids, displayed significantly higher auxin levels than those detected in the *auxT* mutant (**Figure 5**). This supports the notion that AuxT contributes to the accumulation or distribution of these phytohormones inside the nodule. Given the central role of auxins in regulating nodule organogenesis and cell differentiation during symbiosis, it is plausible that bEV-mediated auxin transport constitutes an additional mechanism through which rhizobia modulate nodule development (Kohlen *et al*., 2018). In this context, AuxT could promote the incorporation or release of auxins via bEVs, thereby contributing to the local regulation of hormonal balance within the symbiosome and, ultimately, to the efficiency of the symbiotic interaction. In this sense, the symbiotic impact of AuxT on the *S. fredii* HH103-*G. max* interaction was demonstrated; plants inoculated with the *auxT* mutant exhibited a significant reduction in symbiotic parameters (**Figure 6**). The decrease in both nodule number and biomass, as well as shoot growth, indicates that the absence of this transporter impairs the establishment of symbiosis or optimal nodule function and development. Crucially, the complemented strain restored the wild-type phenotype, confirming that this effect stems specifically from *auxT* deletion (**Figure 6**). Although the underlying molecular mechanism cannot be established solely from these assays, these results align with the potential involvement of AuxT in auxin transport during rhizobial colonization.

Collectively, these results allow us to propose a model where AuxT functions as a bacterial transporter involved in mobilizing auxins produced by *S. fredii* HH103 into the periplasmic space and ultimately into bEVs (**Figure 7**).

**Figure 7.**
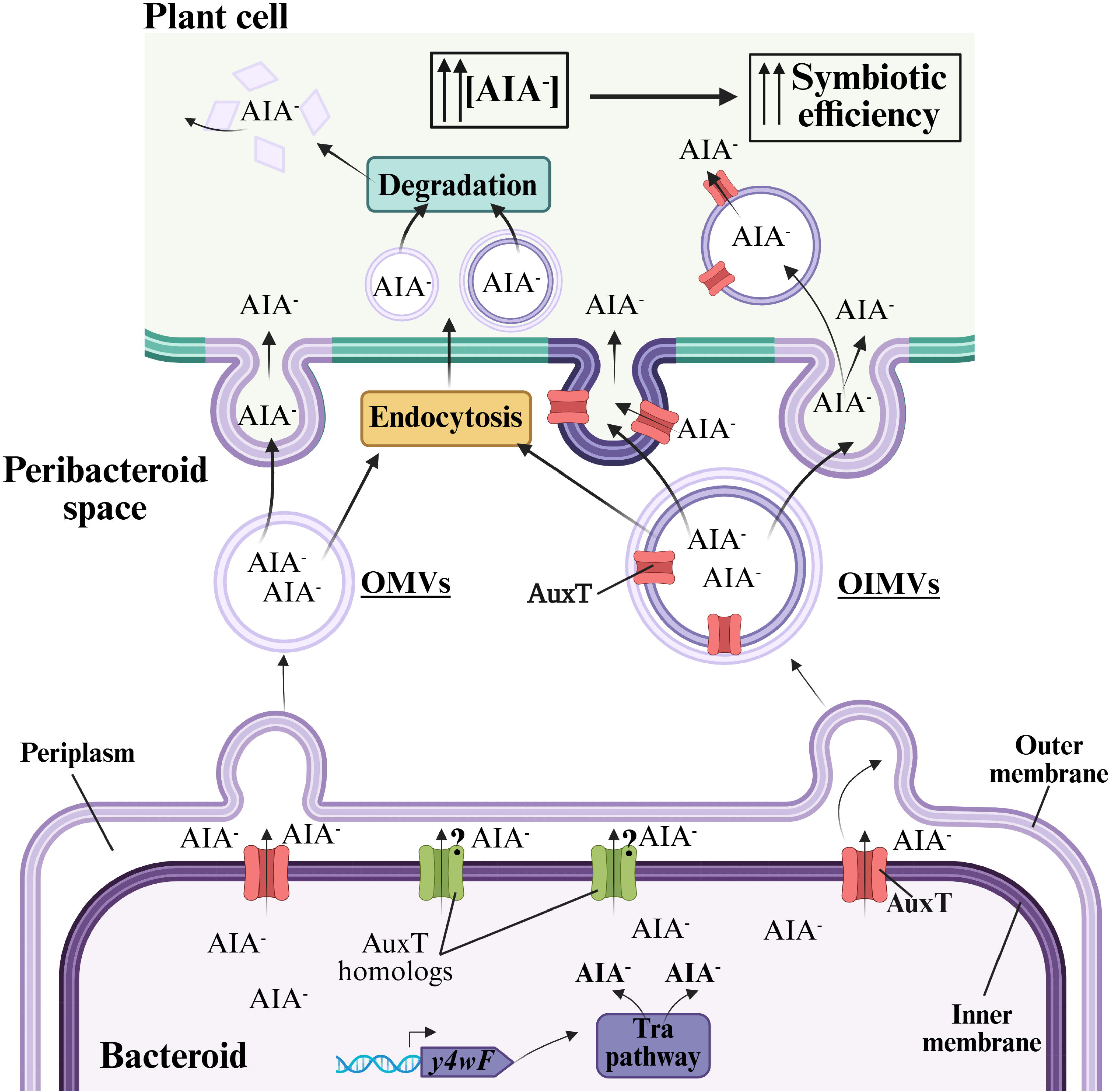
Schematic model of auxin synthesis, translocation, packaging into bEVs and delivery to the host plant by *S. fredii* HH103 during late-stage symbiosis. Auxins synthesized in bacteroids through the constitutive Tra pathway are proposed to be transported from the cytoplasm to the periplasm by the inner membrane transporter AuxT (red), likely in cooperation with its homologous transporters (green). Once in the periplasm, auxins are proposed to be loaded into outer membrane vesicles (probably B-OMVs) and/or outer-inner membrane vesicles (likely B-OIMVs). In OIMVs, AuxT is proposed to remain associated with the inner membrane. After crossing the peribacteroid space, bEVs may deliver auxins to the host cell either by direct membrane fusion or by endocytic uptake followed by vesicle degradation. Membrane fusion would result in the direct release of auxins into the plant cytoplasm and, in the case of OIMVs, could also incorporate the inner membrane containing AuxT into the host membrane. Alternatively, endocytosed bEVs would release their auxin cargo following vesicle degradation, increasing intracellular auxin levels in the host cell. The resulting elevation in auxin concentration is proposed to enhance symbiotic efficiency by promoting optimal nodule function and plant growth. Figure created with BioRender.

According to this model, auxin production in *S. fredii* HH103 is driven primarily by the constitutive Tra pathway associated with the *y4wF* gene, maintaining a sustained basal but high synthesis of these phytohormones (Costacurta and Vanderleyden 1995; Patten and Glick 1996). Cytoplasmic auxins are subsequently translocated to the periplasm by AuxT located in the inner membrane, likely in coordination with its homologous transporters. Once in the periplasm, auxins are incorporated into bEVs generated through distinct vesiculation mechanisms, predominantly B-OMVs and B-OIMVs release. Notably, explosive cell lysis is unlikely to contribute substantially to bEV biogenesis in bacteroids, as ribosomal proteins were completely absent from bacteroid-derived bEVs, suggesting that lysis-derived vesicles are not a major bEV population (Ayala-García *et al*., 2024a). Once released, auxin-loaded bEVs could transfer the phytohormones to plant cells through distinct mechanisms depending on their vesicle type and physiological context. OMVs could fuse with the peribacteroid membrane, discharging their auxin content directly into the plant cell cytoplasm, or they could be internalized via endocytosis and subsequently degraded, leading to auxin release. Alternatively, OIMVs (which harbor both inner and outer membranes) could likewise undergo endocytosis and vacuolar degradation to release encapsulated auxins. Another intriguing possibility is that this kind of bEVs fuse with the peribacteroid membrane via their outer membrane, delivering the vesicular lumen alongside the inner membrane and intermembrane molecules into the plant cytoplasm, which contains the auxins. In all these scenarios, the result would be a local increase in auxin concentration within the host plant cell. While these pathways provide a robust framework for bacteroid bEV-host interactions, the mechanisms operating during other symbiotic stages remain uncharacterized. Elevated auxin levels inside the nodule have been associated not only with changes in plant physiology but also with a profound reprogramming of bacterial metabolism, increasing the expression of nitrogen fixation and energy metabolism genes in bacteroids, as well as nitrogenase activity and overall symbiotic efficiency (Defez *et al*., 2019).

In summary, this work establishes a comprehensive model for bEV-mediated phytohormone delivery during the *S. fredii* HH103-*G. max* symbiosis. Through the functional deployment of AuxT, rhizobia are able to actively compartmentalize and translocate intracellularly synthesized auxins into bEVs. The severe symbiotic impairment observed in the *auxT* mutant underscores that this transport mechanism is indispensable for optimal nodule functionality and overall plant growth. Collectively, these results broaden our insight into the rhizobia-legume dialogue, highlighting bEVs as crucial conveyors for modulating host development via phytohormone delivery.

## Conclusion

This study uncovers a previously unrecognized mechanism of interdomain communication in which *S. fredii* HH103 uses bEVs to actively remodel the hormonal environment of its host plant. We identify AuxT as the first bacterial PIN-like auxin transporter functionally linked to bEV-mediated phytohormone delivery. Although belonging to the bacterial BART transporter superfamily, AuxT displays remarkable structural convergence with eukaryotic PIN-family auxin transporters and operates independently of the classical flavonoid-induced symbiotic program, enabling the constitutive mobilization of bacterial synthesized auxins for selective loading into bEVs. The severe reductions in nodulation, nodule biomass, and plant growth observed in the *auxT* mutant demonstrate that this bEV system-mediated hormone transport pathway is essential for maximizing symbiotic efficiency.

Beyond establishing a new paradigm for bEV function, our findings reveal that rhizobia employ dedicated membrane transport systems not only to translocate phytohormones across bacterial compartments but also for their fine-tune delivery across the symbiotic interface, thereby directly influencing host developmental progress. This work expands our understanding of how bacteria manipulate plant signalling through bEV-mediated cargo selection and identifies AuxT as a key determinant of efficient nitrogen-fixing symbiosis.

Given the central role of rhizobium-legume interactions in sustainable agriculture, these discoveries also provide a conceptual framework for developing rhizobial bEV-based bioinoculants with enhanced plant growth-promoting capacity. Engineering strains with optimized AuxT-mediated auxin delivery -or exploiting bEVs as natural nanocarriers for targeted delivery of beneficial molecules- offers promising opportunities to improve biological nitrogen fixation, reduce dependence on synthetic fertilizers, and enhance crop productivity in an environmentally sustainable manner. Together, our findings establish bEV-mediated phytohormone transport as a fundamental mechanism of plant-microbe communication with broad implications for bEV biology, evolutionary cell biology, and agricultural biotechnology.

## Supplementary data

Supplementary data associated with this article can be found in the online version.

## Acknowledgements

We are grateful to Ina Brentrop (Helmholtz Centre for Infection Research, Braunschweig, Germany) for the technical support in the electron microscopy experiments.

## Author contributions

**Natalia Moreno-de Castro:** Validation, Investigation, Visualization. **Mustafa Safa Karagöz:** Validation, Investigation, Visualization. **Irene Herrero Gómez:** Investigation.

**Susanne Sievers:** Methodology, Visualization. **Mathias Müsken:** Methodology, Visualization. **Joaquín Giner-Lamia**: Investigation, Visualization, Review. **Irene Jiménez-Guerrero**: Writing – Review & Editing, Funding Acquisition. **Francisco Pérez-Montaño**: Conceptualization, Writing – Original Draft Preparation, Visualization, Supervision. **José Manuel Borrero-de Acuña**: Conceptualization, Writing – Review & Editing, Project Administration, Funding Acquisition, Supervision.

## Conflict of interest

The manuscript has neither been previously published nor is it under consideration by any other journal. Before submitting this manuscript, we have obtained the consent of all the authors. In addition, our manuscript and research have no conflict of interest.

## Funding

N.M.-C. received the FPU fellowship FPU20/06902 funded by the Spanish Agencia Estatal de Investigación (MCIN/AEI/10.13039/501100011033). This work was supported by research grants: EMERGIA20_00048 and ProyExcel_00450 from the Junta de Andalucía, the PID2021-122395OA-I00, PID2024-156076NB-I00 and TED2021-130357B-I00 from the Spanish Agencia Estatal de Investigación (MCIN/AEI/10.13039/501100011033).

## Data availability

All data reported in this paper will be shared by the lead contact upon request. This paper does not report original code. Any additional information required to reanalyse the data reported in this paper is available from the lead contact upon request.

---

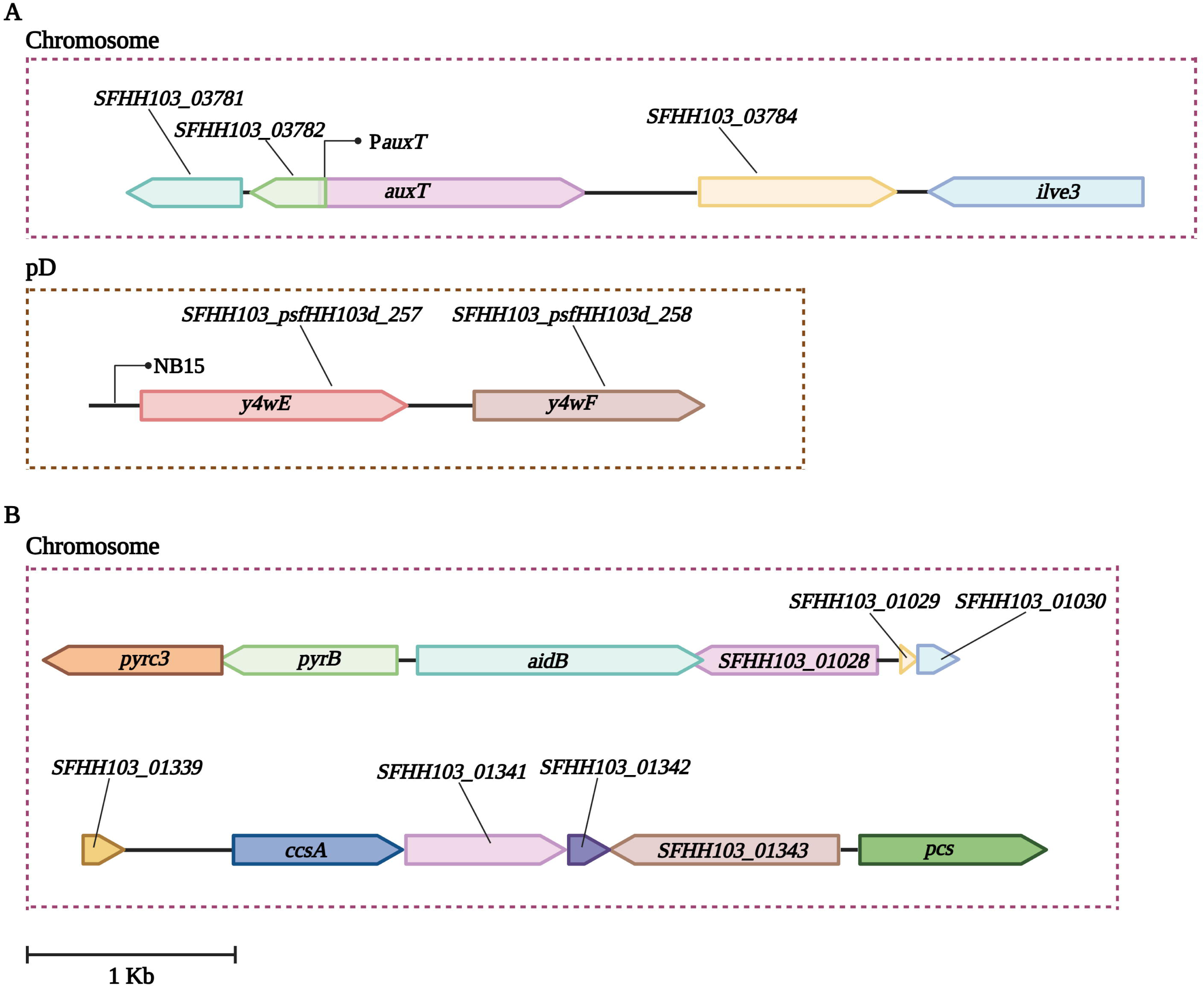

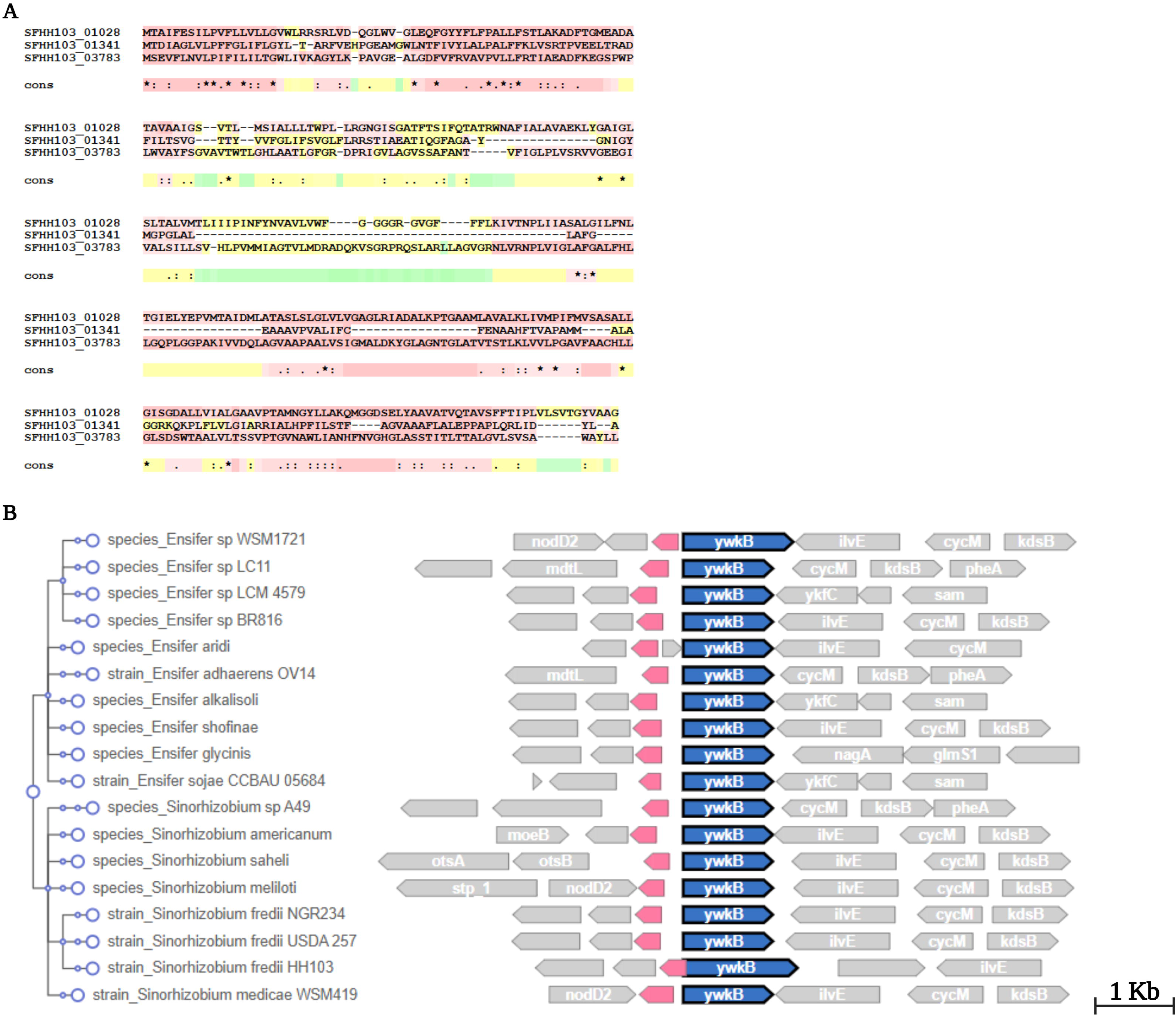

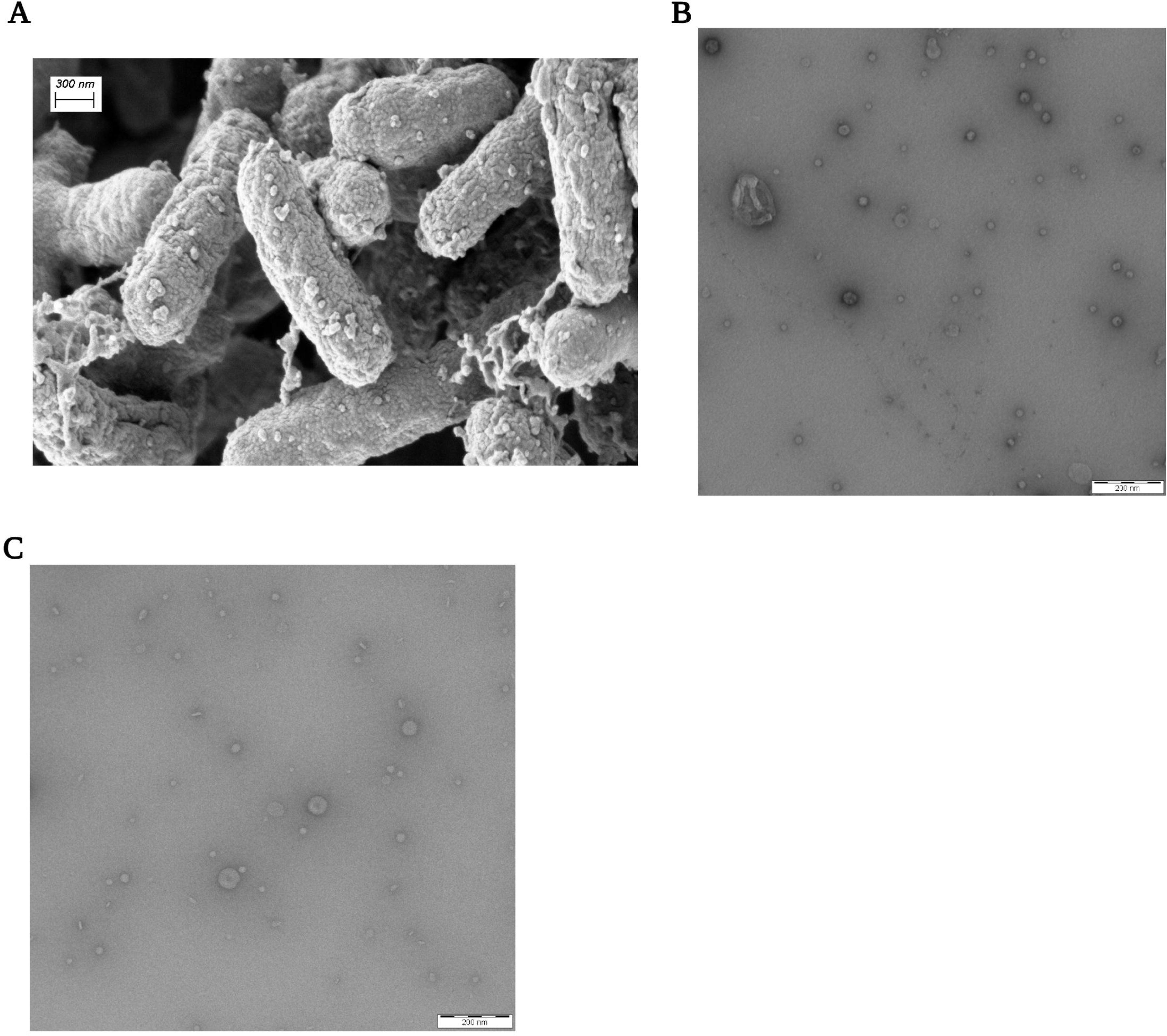

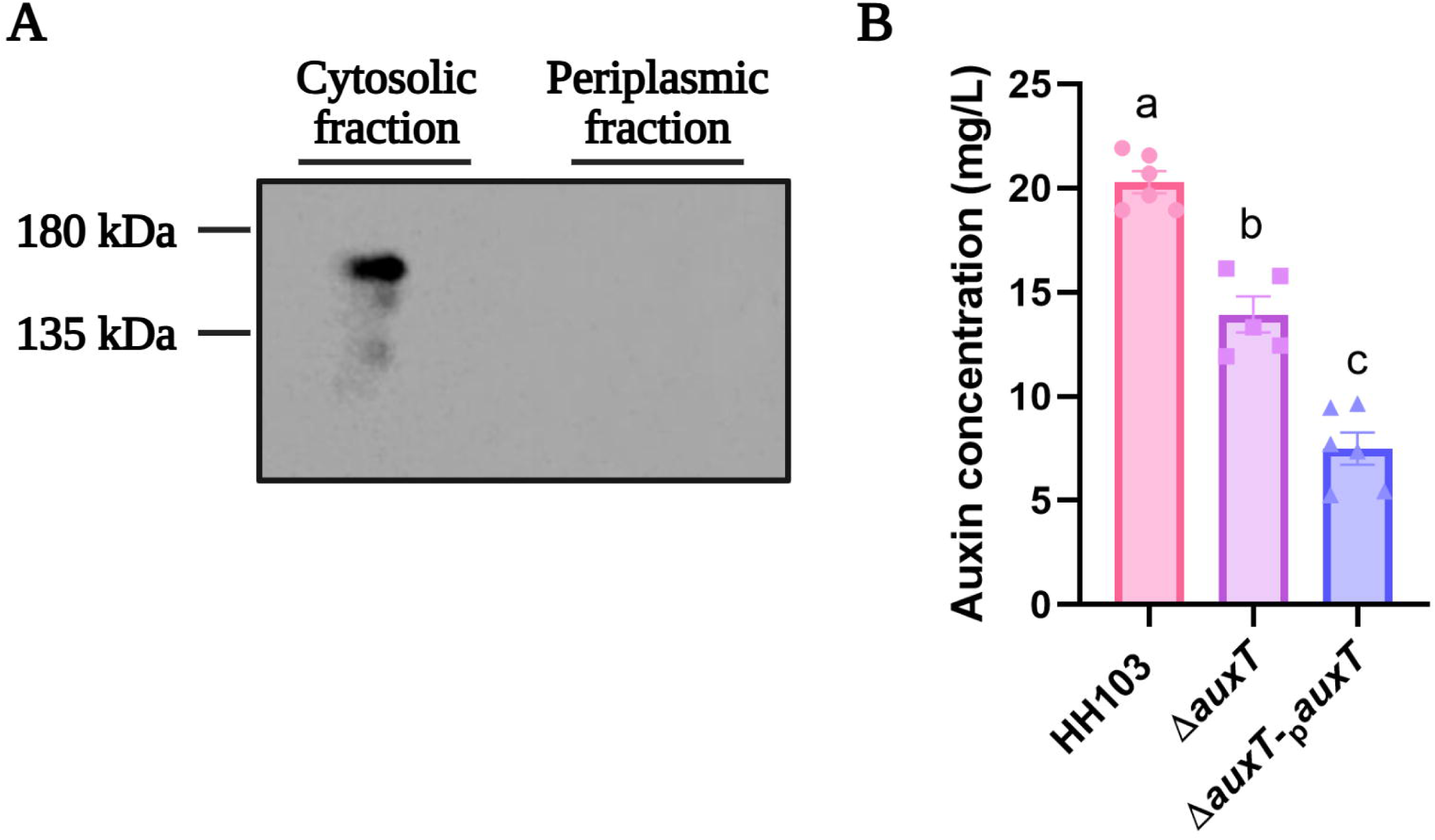

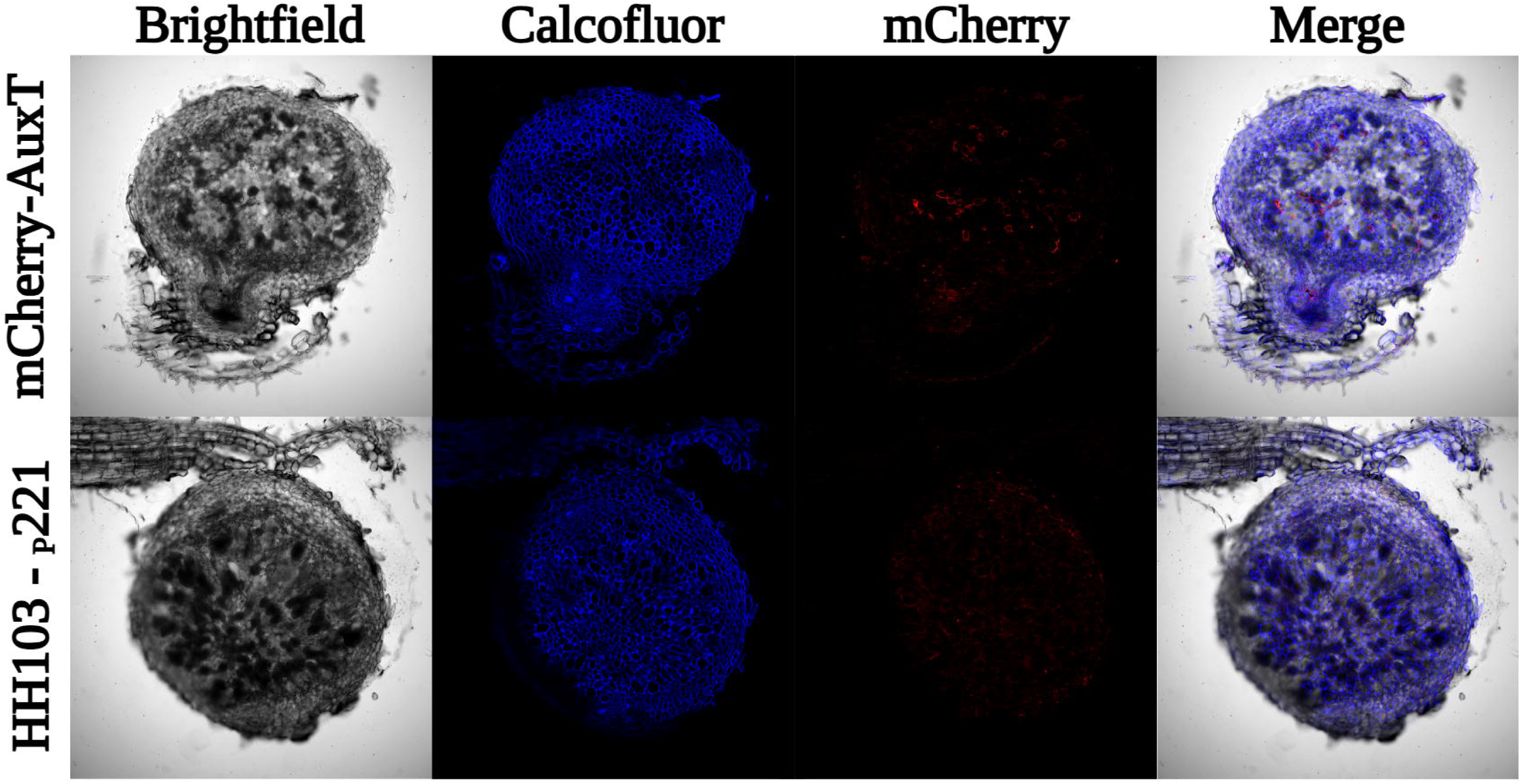

## Notes

### Competing Interest Statement

The authors have declared no competing interest.

